# Transcriptional and Histone acetylation changes associated with CRE elements expose key factors governing the regulatory circuit in early stage of Huntington’s disease models

**DOI:** 10.1101/2023.01.19.524732

**Authors:** Sandra Arancibia-Opazo, J. Sebastián Contreras-Riquelme, Mario Sánchez, Marisol Cisternas-Olmedo, René L. Vidal, Alberto J. M. Martin, Mauricio A. Sáez

## Abstract

Huntington’s disease (HD) is a disorder caused by an abnormal expansion of trinucleotide CAG repeats within the huntingtin (Htt) gene. Under normal conditions, the CREB Binding Protein interacts with CREB elements and acetylates Lysine 27 of Histone 3 to direct the expression of several genes. However, mutant Htt causes depletion of CBP which in turn induces altered histone acetylation patterns and transcriptional deregulation.

Here, we have studied differential expression analysis and H3K27ac variation in 4- and 6-week-old R6/2 mice as a model of juvenile HD. Analysis of differential gene expression and acetylation levels were integrated into Gene Regulatory Networks revealing key regulators involved in the altered transcription cascade.

Our results show changes in acetylation and gene expression levels that are related to impaired neuronal development and key regulators clearly defined in 6-week-old mice are proposed to drive the downstream regulatory cascade in HD.

Here we describe the first approach to determine the relationship among epigenetic changes in the early stages of HD. We determined the existence of changes in pre-symptomatic stages of HD, a starting point for early onset indicators of the progression of this disease.

## Introduction

Within genetic pathologies, there is a group of diseases that involve non-traditional inheritance mechanisms characterized by highly unstable mutations called expansions (McCandless & Cassidy, 2006). Most of these expansions correspond to trinucleotide repeats that include CGG, CAG, CTG, CGG, and GAA, among others. Notably, there are at least 10 neurodegenerative diseases known to be caused by CAG repeat expansions, also referred to as polyglutamine (polyQ) diseases, among which there is Huntington’s disease (HD) (Paulson, 2018; Gonzalez-Alegre, 2019).

HD is an autosomal dominant neurodegenerative disorder caused by the expansion of the PolyQ segment equal to or greater than 36 triplet repeats in exon 1 of the huntingtin gene (*Htt*), thereby producing a polyglutamine tract in the Huntingtin protein (Nopoulos, 2016; Sathasivam et al., 2013; Saudou & Humbert, 2016; Zuccato et al., 2010). HD is characterized by neuronal degeneration and loss of brain striatal tissue (McCandless & Cassidy, 2006; Saudou & Humbert, 2016) due to selective loss of GABAergic projection medium spiny neurons, which represent approximately 90-95% of the neurons present in the striatal region (Gatto et al., 2020; Zheng & Kozloski, 2017). The number of CAG repetitions is associated with the disease penetrance and the appearance of the first symptoms, which in most cases begin between the ages of 30 and 50 years (Bhattacharyya, 2016; Cohen-Carmon & Meshorer, 2012; Landles & Bates, 2004; Schilling et al., 1995). The expansion of 60 or more CAG segment repeats in the Htt gene results in juvenile HD, which usually develops under the age of 20 (Ament et al., 2017; Bossy-Wetzel et al., 2008; Cohen-Carmon & Meshorer, 2012; Gatto et al., 2020; Rubinsztein et al., 1996; Shin et al., 2017). Characteristic HD symptoms include progressive involuntary movements, and behavioral, cognitive, and neuropsychiatric symptoms among other manifestations (Bossy-Wetzel et al., 2008; Gatto et al., 2020; Jimenez-Sanchez et al., 2017; Wang et al., 2018).

To date, the precise cellular function of HTT protein, encoded by the *Htt* gene, has not been yet elucidated (Reinius et al., 2015; Saudou & Humbert, 2016; Schulte & Littleton, 2011), although its role in intracellular trafficking, membrane recycling, neuronal transport, and postsynaptic signaling has been studied (Saudou & Humbert, 2016; Schulte & Littleton, 2011). Mutant Huntingtin protein (mHTT) form is associated with increased mitochondrial dysfunction, alterations in energy metabolism, ER stress and abnormal protein-protein interactions (Kim et al., 2011), all of which can result in dysregulation of transcriptional machinery and altered gene expression pattern into the nucleous (Kim et al., 2011). Several key proteins from the transcriptional machinery aggregated in the presence of mHTT are affected in the processes in which they participate as well as in their function, such proteins include the CREB binding protein (CBP), the specificity protein 1 (SP1), and the TATA-binding protein (TBP), among others, which finally produce an effect over gene transcription (Landles & Bates, 2004).

CBP is a ubiquitously expressed protein that possess acetyltransferase activity and acts as a transcriptional coactivator of several transcription factors (TFs), such as the cAMP response element-binding protein (CREB) (Dai et al., 1996; Henry et al., 2013; Sanchez-Mut & Gräff, 2015). CREB localizes in the cell nucleus where it binds to the cAMP response element (CRE) on the promoter region of several target genes (Wang et al., 2018). To initiate the transcription process, activation of CREB is initiated by phosphorylation of its Serine 133 followed by the recruitment of CBP and posterior stimulation/inhibition of the expression of downstream genes *ATF-3*, *STAT3*, and *BDNF* (Wang et al., 2018). These downstream genes participate in different neural functions such as, stress response (Han et al., 2018), a modulator of synaptic plasticity and synaptic communication (Sasi et al., 2017) among other processes.

Additionally, CBP can activate gene transcription through its intrinsic histone acetyltransferase activity (Miller et al., 2012), adding an acetyl group to lysine residues (K8) of Histone 4 (H4) or H3K27 (H3K27ac), which regulates DNA accessibility to the transcriptional machinery (Dancy & Cole, 2015; McManus & Hendzel, 2001; Miller et al., 2012; Tie et al., 2009; Valor et al., 2011). Studies suggest that abnormal transcriptional regulation plays a role in the development of Huntington’s disease, where CBP sequestration disrupts CRE-mediated transcription (Nucifora et al., 2001; Landles & Bates, 2004).

Although the genetics of HD has been widely studied, most of the results are based on late stage of HD progression models of the disease (Achour et al., 2015; Ferah et al., 2019). Moreover, genomics and epigenomics studies on early stages has been mainly focused on physiological effects such as neurogenesis (Kohl et al., 2010), retinal adaptation to light and dark (Ragauskas et al., 2014), and neuroprotective effects of synaptic modulation (Stack et al., 2007). In order to understand the epigenetic modification in early stage of HD we have used the juvenile HD model to study changes in H3K27 acetylation and their relationship with transcriptional processes in striatal neurons revealing key factors that govern the transcriptional landscape in the early stages of HD.

## Results

### Protein aggregation and transcriptional changes in Huntington’s disease are detectable in pre-symptomatic stages

The alteration of gene expression in striatal neurons is an early event described in HD that starts before neuronal death, being nuclear and cytoplasmatic mHtt inclusion. However, the gene expression profile of the R6/2 HD model before the striatal neuronal loss is not yet described. We first see the presence of mHtt accumulation in striatum tissue at 4, 6, and 13 weeks in the R6/2 model. We observed a progressive increase in mHtt aggregation levels in the striatum and cortex brain region (Figure 1). These results confirm the presence of mHtt before the onset of motor symptoms.

**Figure 1.**
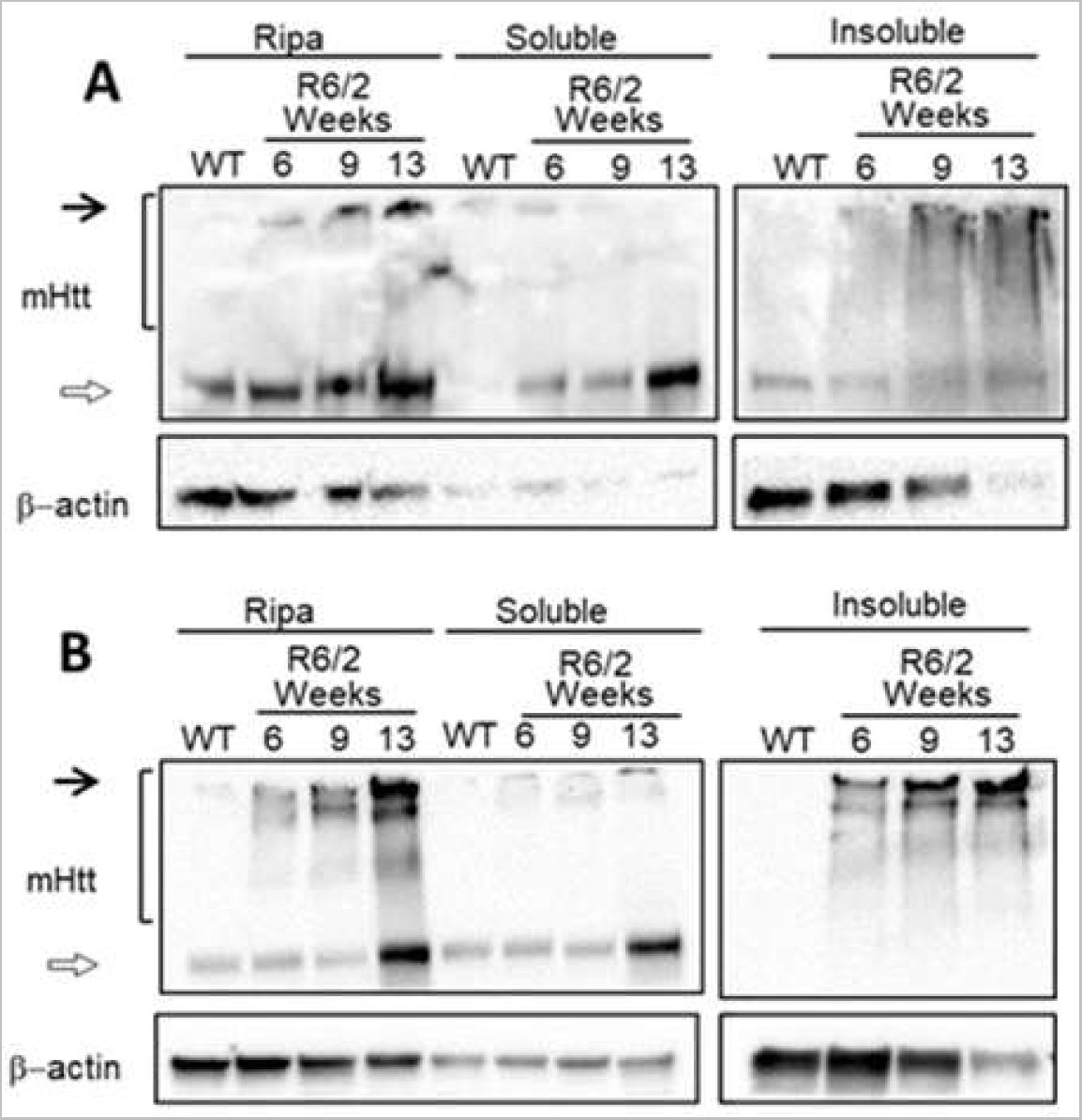
Mutant huntingtin progressively aggregates since 6 weeks. Total protein extraction was performed, using RIPA buffer or separating soluble and insoluble fractions from R6/2 mice (6, 9 and 13 weeks old) and WT mice. **A)** Western blot of striatal brain tissue from WT or R6/2 mice, black arrows indicate high molecular weight species, bracket show smearing of mutant huntingtin. **B)** Western blot from brain cortex tissue from WT or R6/2 mice, black arrows indicate high molecular weight species and bracket show smearing of mutant.

To determine the presence of transcriptomic alterations in the early stages of HD, we performed a differential gene expression analysis. To do so, striatal tissue from both wild type (WT) and R6/2 mice models at 4 and 6 weeks were analyzed. We found that 23 differentially expressed (DE) genes in 4-week-old when comparing WT mice against the R6/2 mice (Figure 2A and Material Supplementary 1), while 1365 DE genes were observed at 6-week-old mice (Figure 2B and Material Supplementary 1). Comparing the DE genes of both ages, we found that there are three genes (*Egr1*, *Rec8*, and *Gm12695*) downregulated in the HD models (see supplementary figure 1).

**Figure 2:**
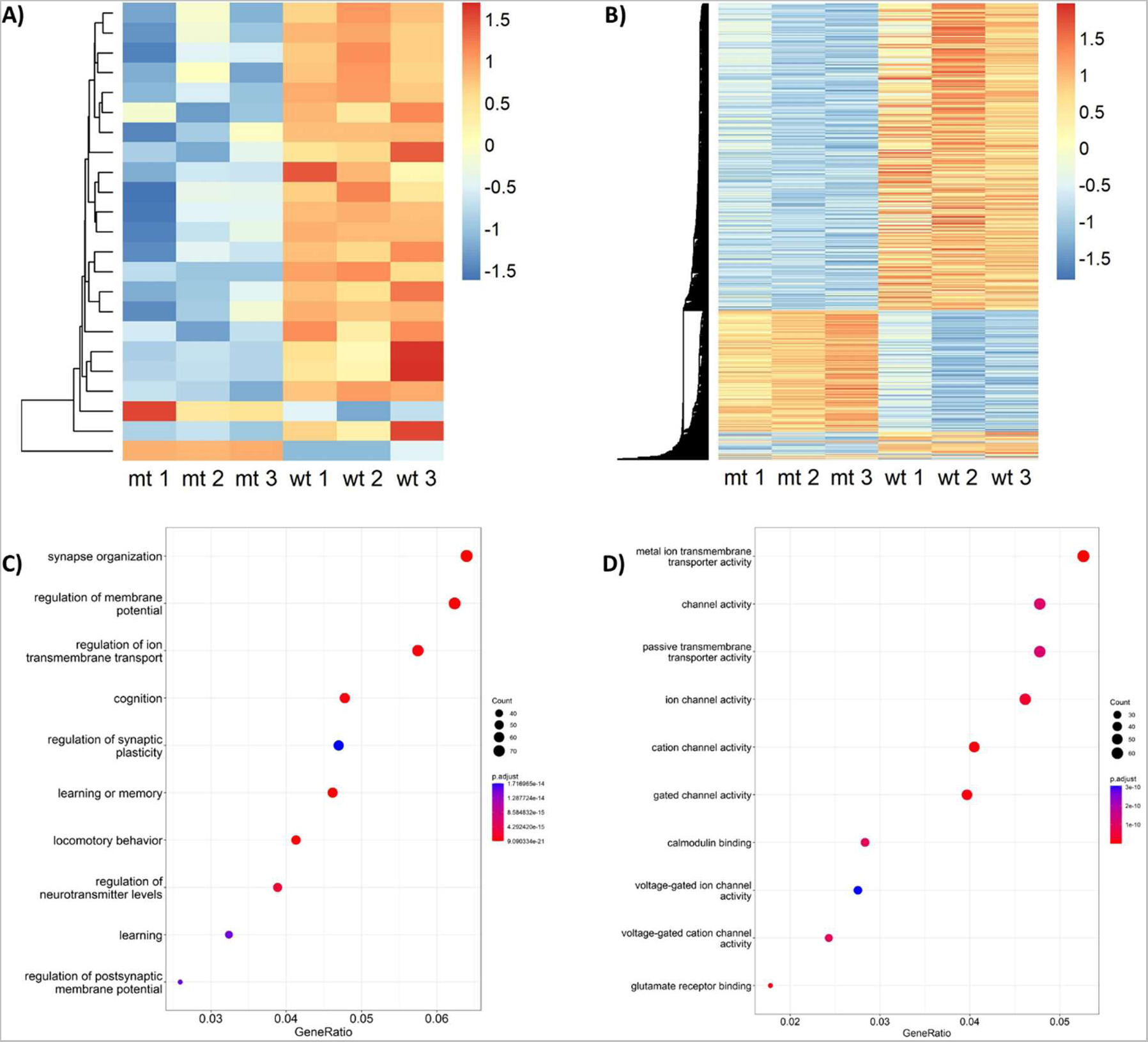
Heatmap of differentially expressed gene levels and their functions found in the R6/2 murine model. A) Heatmap of DE genes in 4-week-old mice. Each row represents one gene, and each column represents one sample. Gene quantification (expressed as z-score) is represented in a color scale ranging from blue (low) to red (high) expressed genes. B) Expression of the changed genes in the R6/2 model at the 6-week control group. C) Top 10 enriched biological processes and D) top 10 molecular functions enriched in the 6-week-old R6/2 mice.

To better understand the DE genes processes, we performed enrichment analysis for metabolic pathways and gene ontology (GO) terms. No significant enrichment for DE genes for metabolic pathways or GO biological processes molecular functions were found at four weeks old. Whereas for six weeks old of R6/2 mice presented molecular function and biological processes GO terms enriched that are related to cell communication functions that could be associated with neurological events present in the context of HD such as synapse organization, cognition, learning and locomotory behavior for biological process and channel activity for molecular function (Figure 2C and D, supplementary figure 2).

### Decreased levels of H3K27ac are detectable in pre-symptomatic stages of Huntington’s disease

In order to determine the epigenetic changes in early stage of HD, we performed chromatin immunoprecipitation (ChIP) analysis in striatum tissue samples to determine the level of H3K27ac in the R6/2 mice at 4 and 6 week old to determine the impact of CBP sequestration by mHTT.

Chromatin immunoprecipitation was validated by qPCR in a positive control for acetylation as a GAPDH promoter (Figure 3A). When an anti-IgG antibody was used, no acetylation of the corresponding promoter region was detected at both 4 and 6 week old. However, when using the anti-H3K27ac antibody a variable level of acetylation which was most significant in the 6 week old group.

**Figure 3:**
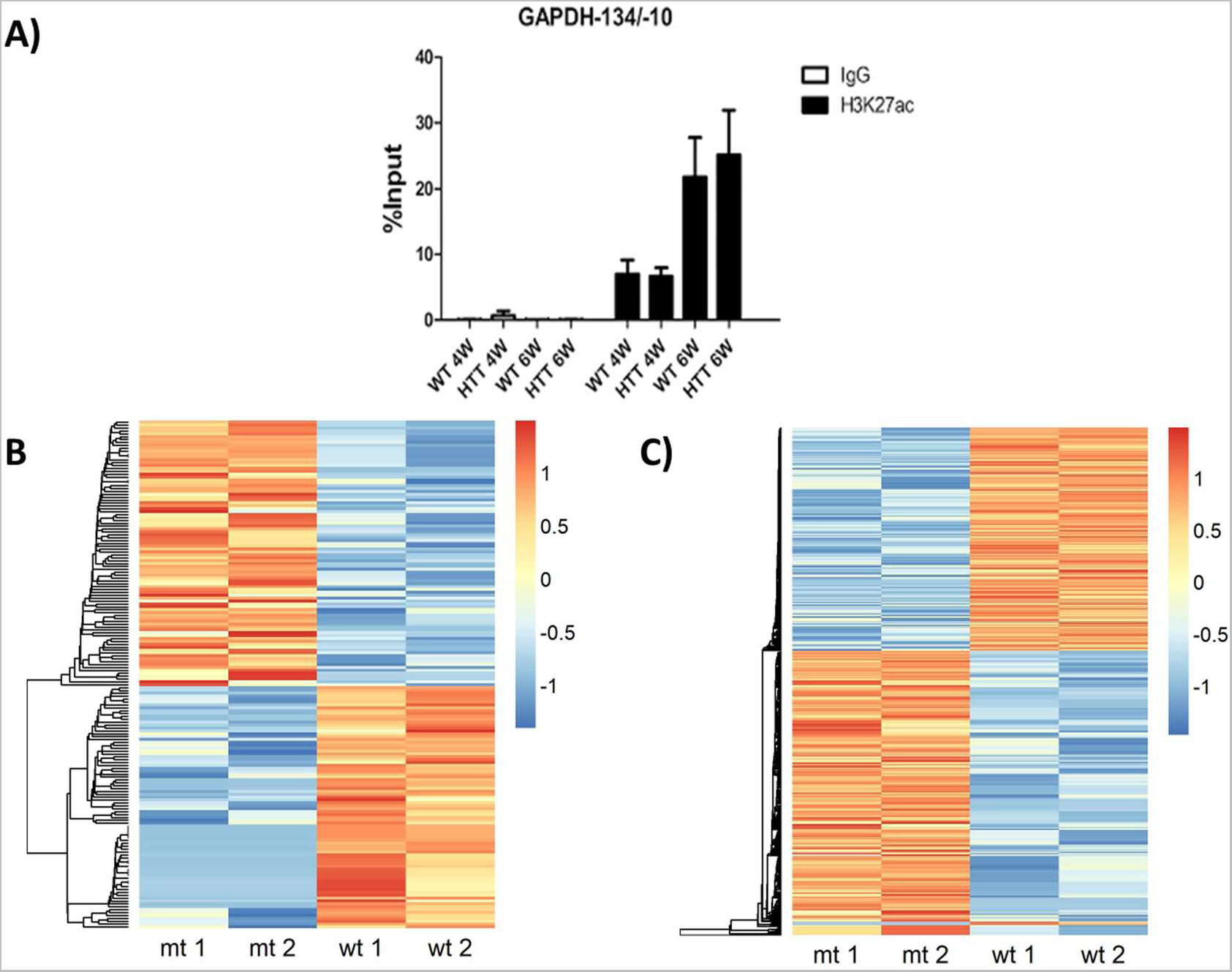
Determination of H3K27 acetylation level in striated tissues in R6/2 and control mice. **A)** qPCR results in the promoter region of GAPDH in 4- and 6-week-old striated tissue samples. Anti-IgG and anti-H3K27ac antibodies were used. **B)** Heatmap representation of promoters with differentially acetylated regions at 4 weeks. **C)** Heatmap representation of promoters with differentially acetylated regions at 6 weeks.

In addition, common H3K27 acetylation peaks were evaluated in control and mutant samples at four and six weeks of age. At 4 week old WT samples showed 129,316 common peaks while the R6/2 samples exhibited 153,605 peaks. Similarly, in the case of the 6 week old WT and R6/2 samples, 126,428 and 146,824 common peaks were observed, respectively (see supplementary figure 4).

Subsequently, the presence of H3K27ac on promoter regions was determined using ChIP-seq data from the murine models under study. 176 differentially acetylated promoter regions were found at 4 weeks old (Figure 3B and Material Supplementary 1), whereas at 6 weeks old, 710 promoter regions were found to be acetylated in this same manner (Figure 3C and Material Supplementary 1).

### CRE-regulated genes present changes in their promoter acetylation levels

We carried out an initial search for genes containing a CRE-binding region to allow a posterior analysis of H3K27ac changes in other CRE-regulated genes. We employed a region of 3Kb upstream and downstream of the transcription start site of each gene to assign CRE-binding sites to their regulation (see supplementary figure 5). We found that 9948 protein-coding genes present at least one CRE-binding site in their 6kb regulatory region. Next, this initial list of genes was curated with publicly available CREB ChIP-seq data, yielding a subset of 1612 genes with a verified CRE-binding site (supplementary figure 6). 80 of these verified genes had been previously described as CREB target genes (see supplementary table 1), among which *Fos*, *Penk*, *Bdnf*, *Gapdh*, *Arc*, and *Stat3* genes have been previously linked to CRE sites (Benito et al., 2011).

In the search for differential acetylation patterns in CRE regions in the R6/2 model, 635 CRE regions were found to be differentially acetylated at 4 week old (Figure 4A and Material Supplementary 1) and 1553 regions at 6 week old (Figure 4B and Material Supplementary 1). Enrichment analysis of metabolic pathways and GO terms of genes whose CRE regions are differentially acetylated in both ages identified pathways related to the regulation of biological activity through phosphorylation of proteins (Figure 4C, Figure 4E, and supp. Figure 7) and in the context of neurogenesis and synapsis (Figure 4D and Figure 4F, Supp. Figure 8).

**Figure 4:**
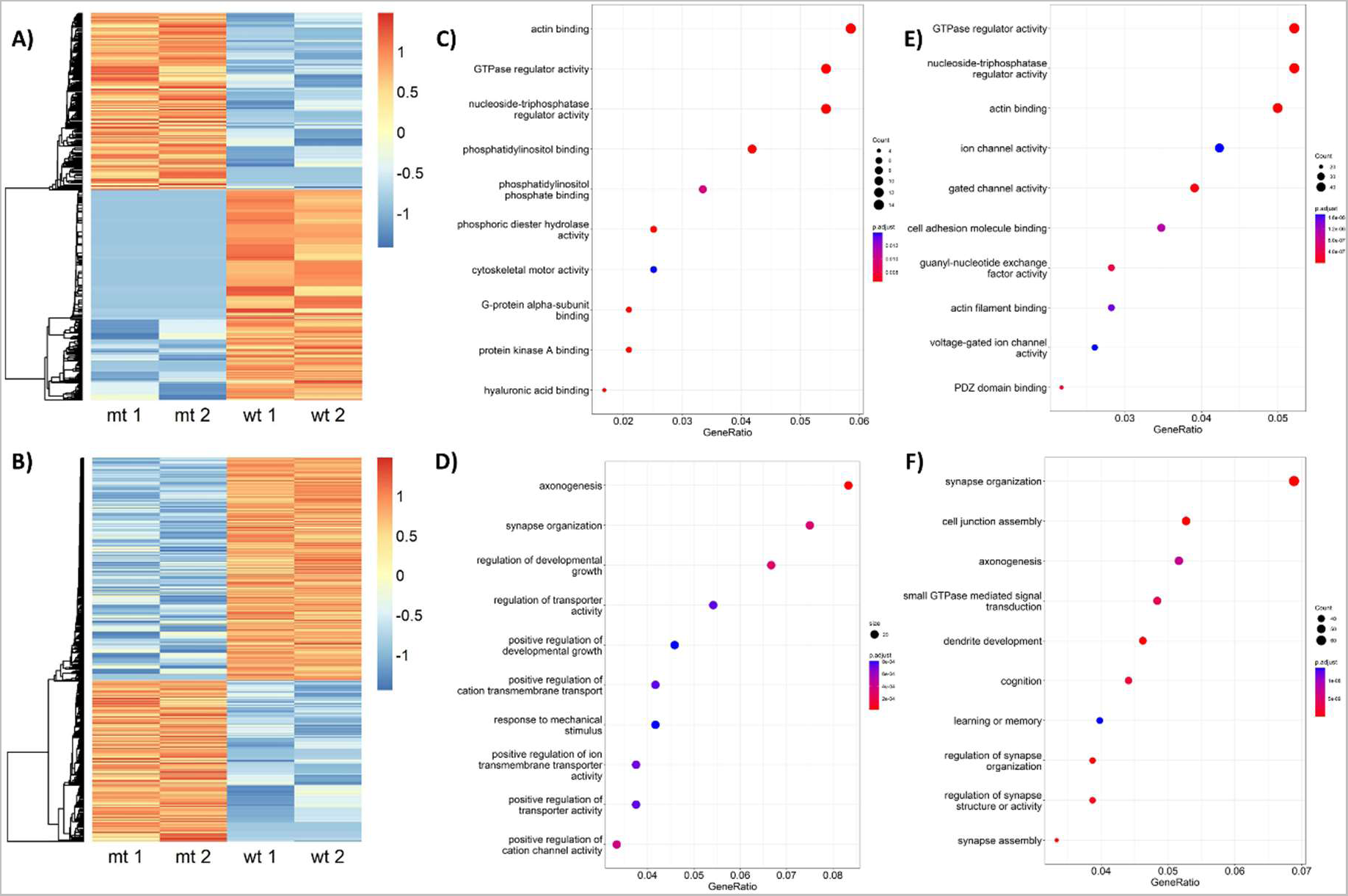
Analysis of CRE element-associated transcriptional changes and their relationship to altered levels of H3K27 acetylation. A) Heatmap of the CRE regions that are differentially acetylated in 4-week-old mice and B) 6-week-old mice. Each row represents the acetylation level (as z-score) from blue (low) to red (high) acetylated regions. C) Top 10 Enrichment of molecular function and D) Top 10 Biological Processes of differentially expressed CRE-region-containing genes observed at four weeks of age. E) Top 10 Enrichment of molecular function and F) Top 10 Biological Processes of differentially expressed CRE-region-containing genes at six weeks of age.

### Altered levels of H3K27ac can be found in differentially expressed genes

The analysis of acetylation over CRE region showed that out of the 635 differentially acetylated CRE regions observed at 4 week old group, 268 are in the gene body, 39 in promoters, two in the gene downstream region and 326 CRE regions are not in the categories above mentioned (Figure 5A). In the case of the 6 week old group, we found a total of 1553 differentially acetylated regions, with 750 placed in the gene body, 188 in promoters, two in the downstream region of the gene, and 613 CRE regions that are not in any of the categories above mentioned (Figure 5B).

**Figure 5:**
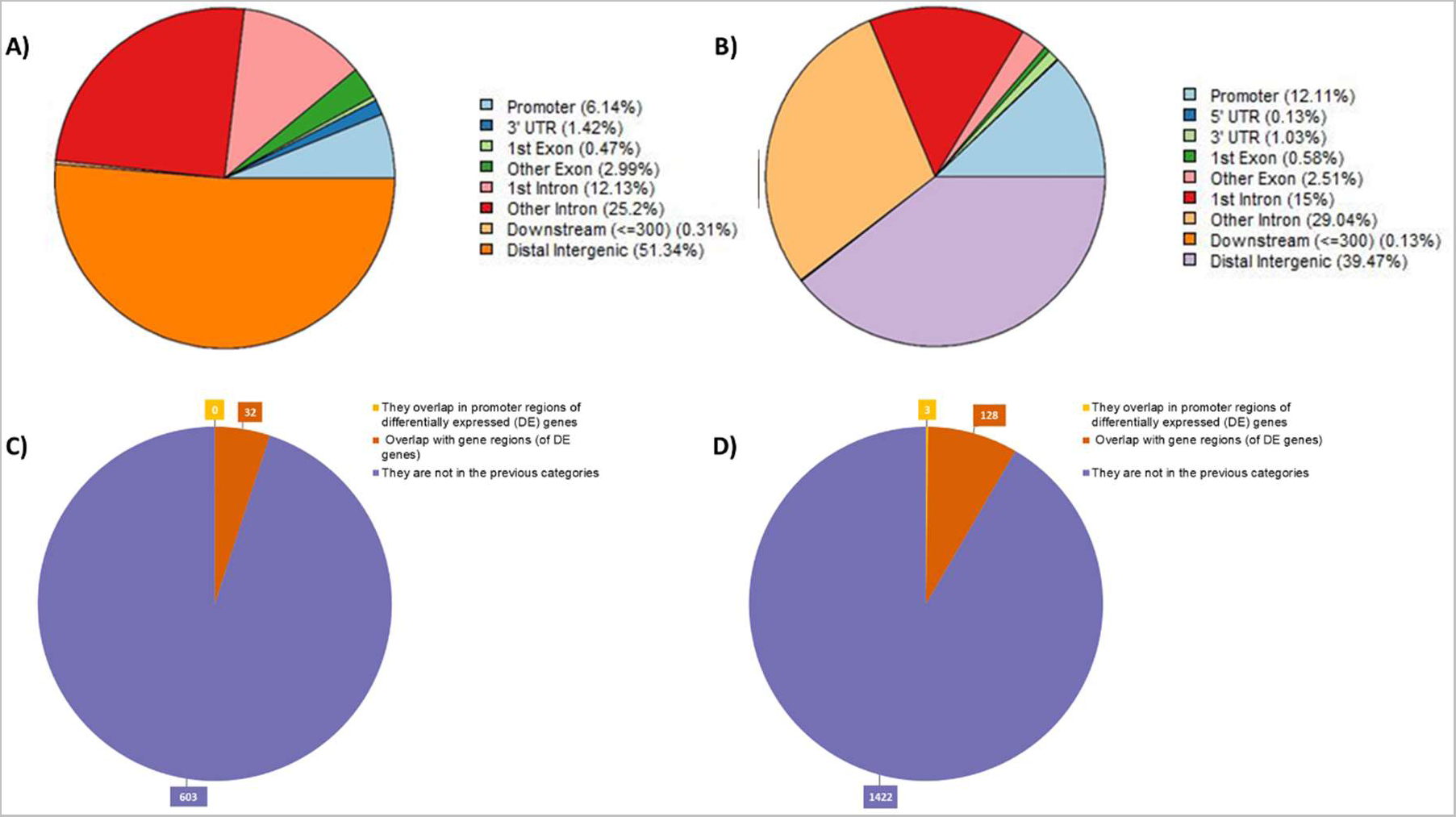
**>A)** Location and relative distribution of differentially acetylated CRE sites at four weeks and **B)** at six weeks of age. **C)** Analysis of differentially acetylated CRE regions located in the promoter or the body of differentially expressed genes in 4-week samples and **D)** in 6-week samples.

Considering these results, we analyzed the differentially acetylated CRE sites in relation to their location in the promoter or body region of the differentially expressed genes. At four weeks of age, from the total number of differentially acetylated CRE sites (635) we found no overlapping between CRE and promoter regions of the differentially expressed genes, 32 CRE regions overlapped with differentially expressed genes, and 603 were not in the above-mentioned categories (Figure 5C). At six weeks of age, three regions overlap with promoter regions of differentially expressed genes, 128 CRE regions overlap with differentially expressed genes, and 1422 are not in the above categories (Figure 5D).

### Changes in the transcriptional landscape in HD related to the CRE-CBP cascade

We next contextualized the general GRN considering the quantification of gene expression carried out in this work for the R 6/2 murine model at four and six weeks of age. For the two ages, both conditions are similar according to the number of nodes and edges presented in the networks, with few elements presented in only one condition (see Table 1).

**Table 1:** Number of nodes and edges (total and unique) in the GRN derived for mutant and control conditions at 4- and 6-weeks old mice.

| Network |  | Total Nodes (TFs) | Unique Nodes (TFs) | Total Connections | Unique Connections |
| --- | --- | --- | --- | --- | --- |
| 4 weeks | Control network | 5876 (643) | 0 | 16635 | 0 |
|  | Mutant network | 5972 (680) | 96 (9) | 17292 | 657 |
| 6 weeks | Control network | 5841 (641) | 1 (1) | 16412 | 22 |
|  | Mutant network | 5925 (663) | 84 (8) | 16990 | 600 |

Looking for a relationship between CRE sites and promoters differentially acetylated and DE genes in 4-week-old mice, we did not find a direct association (Supp. Figure 8). To explore if there is a connection between these genes and the CRE-CREB-CBP cascade, we looked for their first neighbors upstream, in which CREB1 (CREB isoform) and CBP are directing the regulation of some of these genes (Figure 6A and Material Supplementary 1). For the same analysis at 6-weeks old mice, we found changes in the gene expression and acetylation patterns in the promoters related to CREB/CBP, both TFs which are directing changes in the expression of other genes (Figure 6B and Material Supplementary 1). Additionally, several TFs with lower acetylation levels in their promoters direct the expression of several downregulated genes among which *EGR1* and *PDYN* (Figure 6B).

**Figure 6:**
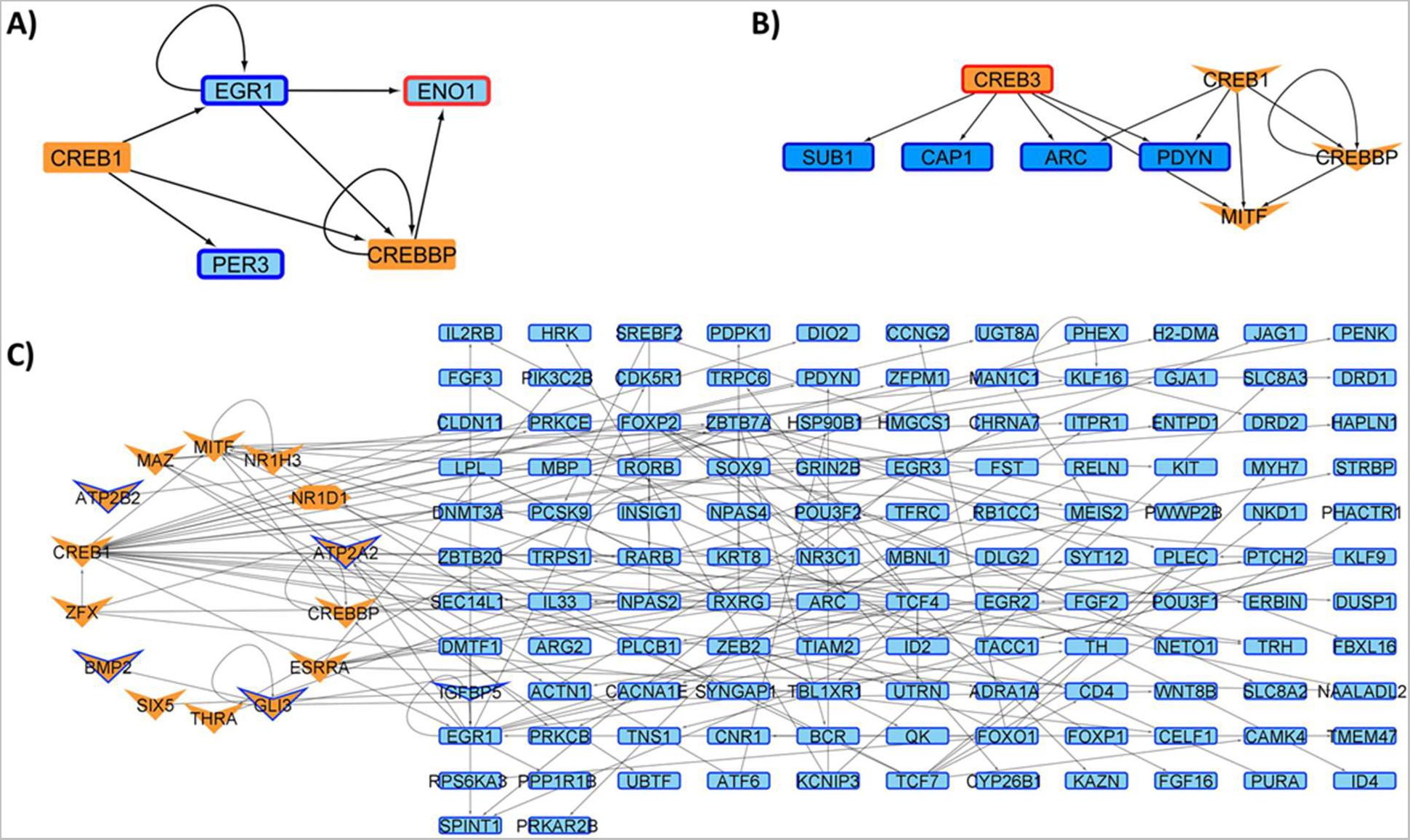
Genes Up (red borders) and down (blue border) regulated by CREB/CBP (orange-colored nodes) in **A)** four weeks-old mice and **B)** Six weeks-old mice. V-shaped nodes represent genes with down acetylated promoters. **C)** Transcription factor coding genes (orange) presenting low H3K27ac levels in their promoter and CRE regions (V-shape and octagon form) directing genes downregulated at six weeks-old Huntington Disease model mice.

### Master regulators guiding the transcriptional landscape in HD presents changes in their expression profiles

Regarding the determination of Master Regulators (MRs), in the four weeks old stage, we did found TF belongs to the MR family or considered as MRs (Supp. Figure 9). Moreover, even if *STAT6* and *EGR1* present a physical interaction, they are not regulated by another MR.

For the 6 weeks networks, several TFs considered MRs due to the dense connectivity between them and physical protein interactions were observed (Figure 6). Within these regulators, 84 are shared for both conditions, while three are defined as MRs only in the HD condition (*Trp63, Pparg* and *Pax6*). Many of these MRs are up/down regulated in the HD condition (Figure 6), with a subset presenting a differentially acetylated promoter region as happens in the CREB isoform CREB1 (Figure 6B and figure 7). Finally, looking for genes that are regulated by these MRs with changes in their expression, we found another TFs and genes with changes in their acetylation levels in promoter/CRE regions (Supp. Figures 10 and 11).

**Figure 7:**
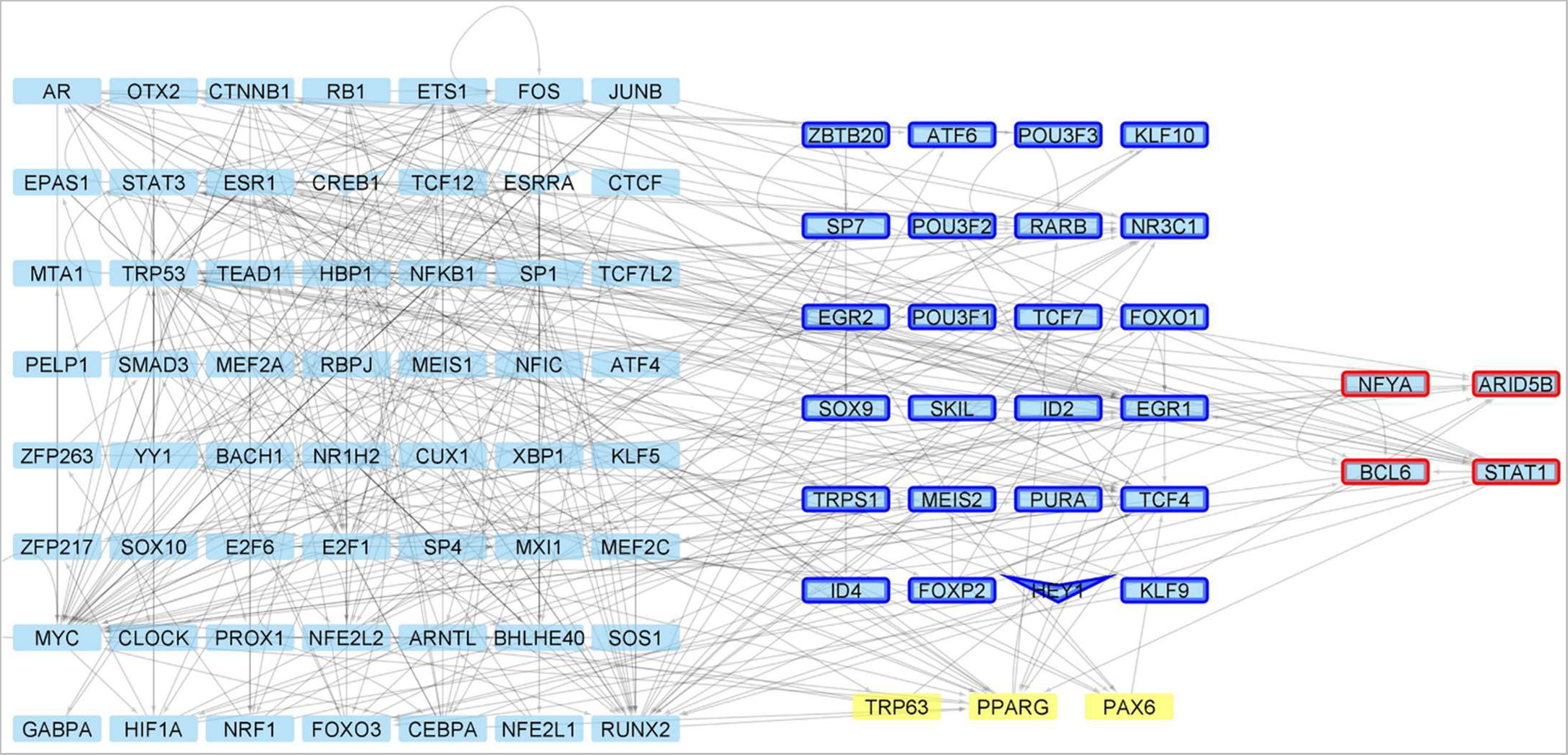
Master Regulators controlling gene expression changes in the HD model at 6 weeks of age. For these MR, there are genes up (red borders) and down (blue border) regulated, with V-shaped nodes representing genes with down acetylated promoters. Yellow TFs are defined as MR only under HD conditions.

## Discussion

Huntington’s disease is neuropathology with no cure and whose treatments focus on improving patient’s quality of life. Current work related to this disease has mainly focused on late stages using the R 6/2 murine model (Achour et al., 2015; Ferah et al., 2019), while studies on early stages have focused, for example, on neurogenesis (Kohl et al., 2010), retinal adaptation to light and dark (Ragauskas et al., 2014) and neuroprotective effects of synaptic modulation (Stack et al., 2007).

In this work, we have addressed the pre-symptomatic stages of Huntington’s disease that have not been addressed in-depth to date in a genomic context, focusing on the integration of data associated with gene expression and changes in H3K27 acetylation levels. We aimed to increase current knowledge on early stages of HD, trying to identify those processes whose functional impairment may serve as an early indicator of the disease. Given that current diagnosis relies on symptom detection or genetic tests that are usually performed when there is the genetic background to do so. Moreover, this search for early markers is also aimed at deciphering key regulatory TFs that are behind these early alterations caused by HD.

Our results show that as time progresses from four to six weeks of age, these mice present an increasing number of differentially expressed genes. Importantly, this increases in the number of DEG indicates that, as expected, younger non symptomatic mice present a little variation when compared to the control mice, but as the mice age increase, the number of DEG increases. Overall, an analysis of the functions and pathways in which these DEGs participate shows enrichment of functional terms linked to ion transport and learning, terms that are related to the functions performed by the striatum, and which are affected as the disease progresses (Rikani et al., 2014; Blumenstock & Dudanova, 2020). At four weeks, we found 21 genes downregulated, which include *Rec8*, *Per3*, and *Egr1*, whereas only two genes are upregulated in HD, *Gm12695* and *Eno1*, which indicates that there is yet a small variation in the cellular functions in the early stages of HD. This trend is also confirmed by the lower number of DEG at this earlier time point than at six weeks. When studying the functions carried out by genes overexpressed in HD at six weeks, we observed that several functions implicitly related to neurodegeneration, linked to genes such as *Adora2a*, *Gm12695* and *Creb3*, are enriched, which somehow is unexpected given that six weeks old mice do not yet show evident symptoms of HD. Furthermore, it is also striking that three genes, *Egr1*, *Rec8* and *Gm12695*, are DE in both ages; both *Egr1* and *Rec8* are downregulated in HD, while *Gm12695* is overexpressed. *Egr1* encodes a TF that participates in transcriptional repression and synaptic plasticity processes (Duclot & Kabbaj, 2017), which are processes known to be impaired in HD. *Rec8* is a protein involved in DNA repair and meiosis, whereas *C1orf87* gene, which is an ortholog of *Gm12695* has been linked to abnormal vocalization and Spinal Muscular Atrophyatrophia^1^. Regarding known DEG genes in HD condition, Crapser and colleagues found a number of genes at eleven weeks, which are downregulated (Crapser et al., 2020).

Among these, the *Gad1* and *Gad2* genes, which encode enzymes that catalyze the synthesis of GABA (Lim et al., 2017), the β4 subunit of the channel MSN voltage-gated sodium *Scn4b* (Miyazaki et al., 2014; Oyama et al., 2006) and *Slc7a11*, which is the astrocyte glutamate transporter, indicate an excitatory/inhibitory balance that is deregulated in this model from early stages of HD (Crapser et al., 2020). In our analysis, *Gad1*, *Gad2*, *Scn4b*, and *Slc7a11*

1. https://www.malacards.org/card/spinal_muscular_atrophy_distal_x_linked_3. genes are down-regulated at 6 weeks of age in the R 6/2 model. This result is consistent with that described by other authors, finding a consistent expression pattern in these genes.

Next, we characterized the occurrence of CRE sites associated with gene regulation of genes. Although there is a database of predicted CREB genes (Xinmin et al., 2005), it is based on an archived version of the genome. For this reason, we have annotated our own catalog of genes putatively possessing a CREB-binding region. This allowed us to perform a functional and pathway-associated analysis of genes whose promoters contain a CREB-binding site that was differentially acetylated. While at four weeks we did not find any metabolic pathways enriched for these CRE regulated genes, many of these genes are ubiquitously expressed, with phenotypes associated to behavior and the nervous system, with functions related to the regulation of transcription (Bult et al., 2019). On the other hand, at six weeks signaling and neurotransmission pathways are enriched on DE genes. This over-representation of genes related to these pathways agrees with previous reports that showed a higher neurotransmitter release in the early stages of the disease. One possible cause for this altered release is the dysfunction and death of MSN striatal neurons, which also is related with the origin of the motor disorders associated with this disease (Bergonzoni et al., 2021). By looking for biological processes in which this subset of CRE-regulated genes is involved, we discovered other functions than those related to neurogenesis. For example, we observed genes such as *Slc5a1, Fgfr2, Col25a1, Fgfr2,* and *Igf* involved in the regulation of transporter activity, synapse organization, axonogenesis, GTPase activator, and regulator activity, functions whose malfunctioning can thus be driving the early stages of the disease. Moreover, the alteration in the expression and regulation of these genes can be due to a possible reduction in striatal neurogenesis originated by the cascade effects triggered by the mutant huntingtin protein, which, in the long run, contributes to the depletion of neurons generated by adults (Bergonzoni et al., 2021; Ernst et al., 2014) and, as a consequence, striatum neurogenesis would be affected (Fedele et al., 2011; Gil-Mohapel et al., 2011), but also of notable relevance is that these processes are related to synapses as described previously. Regardless of the presence of CRE region in genes with changes in their expression levels, there are other sites with changes in their acetylation levels that are not placed in the promoter or the gene body, so further studies must also be focused on how distal regions participate in the juvenile HD.

The integration of information about CRE-binding sites and gene expression levels in the gene regulatory networks shows us that for both time points under study, the transcriptional processes are similar (Table 1). Additionally, although there are no changes in the expression profiles at four weeks for CREB or CBP but given the regulatory networks we generated, it seems that at both ages this CRE-CREB-CBP regulatory cascade would be directly guiding the changes in expression patterns. When looking for TF whose genes have reduced acetylation levels and the downregulated genes controlled by them, we were not able to find a direct relationship at four weeks, probably due to the scarce number of altered genes (Supp. Figure 8), but this is not the case for 6-week-old mice (Figure 6). The regulatory circuit present only in the six weeks time point we studied suggests a relationship between changes in the chromatin status and main affected genes in this context. Comparing these results with other studies, we found genes previously related to HD through differential expression analysis such as *Bcl6*, *Foxp1* and *Egr1* that present changes in their expression profiles, whilst other genes such as Crebbp does not change their expression levels, but present change in their acetylation pattern. These results would indicate that *Bcl6* and *Egr1* could be key regulators whose expression impairment is behind the disease phenotype in early stages of HD (Malla et al., 2021).

Finally, we could not determine master regulators guiding the changes in striatal tissue neurons at four weeks following our approach. This agrees with our findings regarding differential expression and reduced variation on acetylation in the earliest developmental stage we analyzed. However, we have been able to determine several master regulators at six weeks according to the definition proposed by Davis and Rebay (Davis & Rebay, 2017), where although they are mostly shared in the control and disease model condition, several of them are encoded by DEG or show changes in their H3K27 acetylation. It is this variation in expression and acetylation on the MRs genes which could be ultimately guiding the downstream regulatory cascade, which includes genes that are not DE but present promoters that are differentially acetylated (Figure 7 and Supp. Figure 10 and 11). Following this line, within this group of MRs, we also found TFs associated with cell cycle control, which could be associated with neuronal death and other processes that occur in the context of this disease. For example, *Egr1* (Wei et al., 2017), *Foxo1* (Yuan et al., 2014), *Stat1* (Bromberg & Darnell, 2000), *Eno1* (Fu et al., 2015) and *Creb1* (Chae et al., 2015; Fang et al., 2016) are known to be key controllers of cell cycle progression. Among these TFs, the role of *Egr1* should be studied in depth due to its differential expression pattern (downregulated) found in 4- and 6-weeks old mice and its key role as MRs absent in HD we have identified in this work. Furthermore, only three regulators are uniquely defined in the HD model, *Pparg*, *Pax6*, and *Trp63*. *Pparg* and *Pax6* have been previously described as involved in HD and neuronal death (Dickey et al., 2016; Lemprière, 2020), whereas *Trp63* is known to be involved in cell cycle control (Yang et al., 1998) and differentiation of stem cells (Kurita et al., 2004). Importantly, deficiency in *Pparg* has been previously linked to an increase in susceptibility to brain damage (Zhao et al., 2009) and *Pax6* is key for the regulation of neurogenesis (Haubst et al., 2004), linking the overexpression of these two MRs in HD to a corrective response to the characteristic neurodegeneration of HD. Since in the early disease stages we have focused on, mice are still mostly asymptomatic, our findings indicate a role for these genes protecting neural damage in early HD.

This work consists of a first approach to determine the relationship among epigenetics changes in the HD and how it could be affecting transcriptional levels occurring in the early stages of Huntington’s disease. Therefore, further work could validate the role of genes coding for the MR we found as to be related to early HD response as shown in the network analysis. Moreover, genes shown to be linked to these MR and their functions would be a starting point to drive the discovery and determination of possible therapeutic targets and treatments to attack in this disease.

## Materials and Methods

### Mouse strains

Striated tissue samples were obtained from R6/2 mice at their fourth and sixth week of life. The R6/2 transgenic model that contains exon 1 of the mHtt gene human, was obtained from The Jackson Laboratory with all animal protocols abiding the animal management protocol No. 01/2018, approved by the Bioethics and Biosafety Committee of the Universidad Mayor.

### Transcriptomic analysis of striatal tissues from R6/2 mice

Samples of striated tissue from 4- and 6-week-old R6/2 mice and respective controls were obtained in triplicate. Total RNA was purified by tissue homogenization using Trizol® reagent (Invitrogen). The total RNA extraction was digested with DNAse I to avoid contamination of the sample with genomic DNA. The RNA concentration was then measured with the Quant-iT ™ RiboGreen® RNA Assay Kit (Life Technologies), and its integrity was determined by the Bioanalyzer 2100 kit (Agilent Technologies, Santa Clara, CA, USA). RNA libraries were prepared with the Illumina TruSeq TM RNA Sample Preparation Kit, following the manufacturer’s protocol. The rRNA-depleted RNA was fragmented using divalent cations at elevated temperatures. First-strand cDNA synthesis produced single-strand DNA copies from RNA fragmented by reverse transcription. After second-stranded cDNA synthesis, the double-stranded DNA will be repaired at the 3’ ends. Finally, PCR ligated the universal adapters to the cDNA fragments to produce the final sequencing library. After library validation was carried out with the DNA 1000 chip (on the Agilent Technologies 2100 bioanalyzer), the samples were pooled in equal concentrations into one pool and run on the Illumina HiSeq equipment for 100 end sequencing cycles paired.

### Identification of differentially expressed genes in the R6/2 HD mouse model

The RNA-seq Raw data identified so far for striated tissue samples of animal models of HD R 6/2 for 4 and 6 weeks, this data was analyzed using a pipeline explained next: First, the quality of the RNAseq data readings was analyzed through FastQC (Andrews S. 2010); Then, the adapters were removed with Trimmomatic (Bolger et al., 2014); then the sequenced reads were aligned against the mm10 mouse reference genome using STAR (Dobin et al., 2013) and the count of the reads that align against each gene was performed using the Htseq-count (Anders et al., 2015), Finally, the differential expression of the libraries analyzed with the Deseq2 tool (Love et al., 2014).

### ChIP analysis in the R 6/2 model

ChIP tests were performed on cross-linked chromatin samples, with the following modifications. The tissues were washed with a cold 1 × PBS buffer, incubated for 10 minutes with 1% formaldehyde (FA) at room temperature, and washed again with 1 × PBS. For ChIP against chromatin-modifying enzymes, double crosslinking with EGS (ethylene glycol bis (succinic acid N-hydroxysuccinimide ester; E3257; Sigma-Aldrich) was used: FA crosslinked tissue was incubated with EGS for 1 hour at room temperature with gentle agitation, were washed three times with cold PBS, resuspended in 1 ml of cell lysis buffer (5 mM HEPES, pH 8.0, 85 mM KCl, Triton X-100 and proteinase inhibitors) and homogenized with a Dounce homogenizer (Kimble, Vineland, NJ) (~ 25 times using a mortar.) The cell extract was collected by centrifugation at 3,000 g, resuspended in 0.3 ml of sonication buffer (50 mM HEPES, pH 7.9, 140 mM NaCl, EDTA 1 m, 1 % Triton X-100, 0.1 % deoxycholate acid, 0.1 % SDS and a mixture of proteinase inhibitors), and incubate for 10 minutes on ice Extracts from Chromatin sonicated for 15 minutes, 30 seconds, 30 seconds in BioruptorPico (Diagenode) and centrifuged at 16,000 g for 15 min at 4 °C to obtain fragments of 500 bp or smaller. The supernatant was collected, divided into aliquots, frozen in liquid nitrogen, and stored at −80 °C; an aliquot was used for quantification by Qubit fluorometric quantification (Thermo Fischer).

Chromatin size was confirmed by electrophoretic analysis. Extracts of cross-linked chromatin (500 ng) were resuspended in sonication buffer to a final volume of 500 μl; Samples were pre-cleaned by incubating with 2–4 μg of normal immunoglobulin G (IgG) and 50 μl of A / G-agarose protein microspheres (Santa Cruz Biotechnology) for 1 hour at 4 °C with agitation for preclearance. The chromatin was centrifuged at 4,000 g for 5 minutes, and the supernatant was collected and immunoprecipitated with specific antibodies for 12-16 h at 4 ° C. The immune complexes recovered with an additional 50 μl of magnetic beads (Dynabeads, Invitrogen), followed by incubation for 1 hour at 4 °C with gentle agitation. The immunoprecipitated complexes were washed once with sonication buffer, with LiCl buffer (100 mM Tris-HCl, pH 8.0, 500 mM LiCl, 0.1 % Nonidet P40 and 0.1 % deoxycholic acid), and once with Tris-EDTA buffer (TE) pH 8.0 (2 mM EDTA and 50 mM Tris-HCl, pH 8.0), each time for 5 minutes at 4 °C; This was followed by centrifugation at 4,000 g for 5 min. Protein-DNA complexes were eluted by incubation with 100 μl elution buffer (50 mM NaHCO3 and 1 % SDS) for 15 minutes at 65 °C. Extracts were incubated for 12-16 h at 65 °C, to reverse crosslinking the proteins digested with 100 μg / ml proteinase K for 2 hours at 50 °C, and the DNA was recovered by phenol/chloroform extraction and ethanol precipitation using glycogen (20 μg / ml) as the precipitation vehicle. The following antibodies were used for this protocol; anti-H3K27ac antibodies. The following antibodies were used for this protocol; anti-H3K27ac antibodies.

### Determination of mHtt protein levels in brain tissue

The right cerebral hemisphere is dissected into the different areas of the brain to be analyzed. The tissues were immediately transferred to dry ice and stored at −80°C for further analysis at −80°C for further analysis. The brain samples were homogenized in RIPA with 1% SDS and protease inhibitor cocktail (1 tablet per 10 ml, Roche complete, EDTA-free, Sigma-Aldrich) using a sonicator with a 10 s pulse at 40% amplitude.

Protein concentrations were estimated using the BCA assay (Pierce). Cell lysates was denatured under reducing conditions by boiling 30 μg of total protein with 1 M DTT and 4X LDS sample buffer (Invitrogen) at 95°C for 5 min. SDS-PAGE was performed using Precast 4-15% Bis-Tris gels (Bio-Rad). Gels were run in 1X tris-acetate SDS running buffer for 2 hr and transferred to a 0.45 μm PVDF membrane using the Turbo Trans-Blot transfer system (Bio-Rad) for 30 min. Membranes were blocked with 5% skim milk in PBS with 0.1% Tween 20 (PBS-T) for 1 hour at room temperature and incubated with reconstituted anti-HTT (MAB5374, 1:1000) and anti-b-actin HRP (sc-47778 1:5000) primary antibodies in 5% skim milk overnight at 4°C.

### Soluble and Insoluble Huntingtin Species

The brain samples were homogenized in Soluble” lysis buffer (10 mM Tris pH 7.4, 1% Triton-X 100, 150 mM NaCl, 10% glycerol) douncing slowly 30 times in glass 1 mL douncer. The homogenate was transferred into labeled, “Insoluble” tubes and lyse on ice for 1 hour. the samples were centrifuged at 4 °C at 15,000 x g for 20 min. The supernatant was recovered as the “Soluble” fraction. The pellet was washed with 500 μL of lysis buffer (x2) and centrifuge at 4 °C at 15,000 x g for 5 min after each wash. Then, the pellet was resuspended with 150 μL lysis buffer supplemented with 4% SDS and sonicate each sample for 30 second at room temperature with a probe sonicator at 40% of amplitud (Insoluble fraction). Finally, boil the insoluble fraction for 30 min. Protein concentrations were estimated using the BCA assay (Pierce). SDS-PAGE was performed using Precast 4-15% Bis-Tris gels (Bio-Rad). Gels were run in 1X tris-acetate SDS running buffer for 2 hour and transferred to a 0.45 μm PVDF membrane using the Turbo Trans-Blot transfer system (Bio-Rad) for 30 minutes. Membranes were blocked with 5% skim milk in PBS with 0.1% Tween 20 (PBS-T) for 1 hour at room temperature and incubated with reconstituted anti-HTT (MAB5374, 1:1000), and anti-b-actin HRP (sc-47778 1:5000) primary antibodies in 5% skim milk overnight at 4°C.

### Identification of H3K27ac level in R 6/2 murine models

Sequencing results were analyzed using a bioinformatics pipeline to identify changes and/or similarities in H3K27ac levels in striatal tissue as follows. First, the quality of the reads in the ChIP-seq data was analyzed via FastQC (Andrews S. 2010); next, adapters were removed with Trimmomatic (Bolger et al., 2014); genome indices were generated, and alignment of the reads was performed using Bowtie against the mouse genome version mm10. We identified enriched regions and peak annotations were analyzed with MACS3 software using a broad-cutoff of 0.1, the max-gap option of 200, and a p-value of 0.05 (Zhang et al., 2008). The acetylation peaks obtained in each sample and those common to both ages under study were analyzed using Bedtools software using the intersectbed option. The count of the reads that align against each gene was performed using the Htseq-count (Anders et al., 2015) and finally, differential analysis of count data was performed with the Deseq2 tool (Love et al., 2014) considering a pValue value of <0.05.

### Quantification of differentially acetylated promoters

Differentially acetylated promoters were identified following the same steps used to identify H3K27ac levels. However, after alignment of the reads using Bowtie against the mouse genome version mm10, the count of the reads that align against each promoter was performed using the Htseq-count (Anders et al., 2015), and the list of promoters obtained from 1000 up and downs stream of the TSS. Differentially acetylated promoters were detected with the Deseq2 tool (Love et al., 2014) over the generated list considering a p-value of <0.05.

### Determination of CRE regions and the genes associated with these regions in mice

CRE-regulated candidate genes were identified using specific distance thresholds from the transcription start site and the occurrence of CREB-binding motif sequences in the genome. To identify CRE-binding regions in the mouse genome, we first downloaded CREB-binding motifs (position-weighted sets) of CRE elements from the JASPAR database (Khan et al., 2018) scanned in the mm10 version mouse genome for occurrences using FIMO software with default parameters (Grant et al., 2011). Subsequently, by creating a script in R language, we found the putative transcription factor-regulated genes that bind to the CRE element, thus determining the putative transcription factor-regulated genes that bind to the CRE element by assigning the CRE elements to the regulation of those genes that have their transcription start site (TSS) using the distances of 3Kb: +3Kb distance from the TSS.

A literature curation of the method was performed comparing the list of genes regulated by CRE that we collected from the literature to the list of genes predicted by the script. Next, we analyzed CREB ChIP-seq data to check whether the CRE regions determined in the genome had been previously described as CREB-binding. Finally, validation of these motifs was performed using CREB ChIP-seq data (mammary tissue, kidney, and neuron), obtained from a public SRA database.

### Identification of altered regulatory events associated with variation in H3K27 acetylation levels and CRE regions

Given the interaction between CBP with mHTT, we expected a variation in H3K27 acetylation levels, affecting the CRE-regulated transcriptional cascade. For this reason, we considered RNA-seq data generated to map genes expressed in a Gene Regulatory Network. These networks’ changes in the transcriptional cascade due to variation in their acetylation level were analyzed to propose a model of how these alterations might guide HD.

### Generation of gene regulation networks in the R6/2 murine model of HD and their respective Wild Type

Considering the expressed genes in both models (R 6/2 and their controls at 4 and 6 weeks old), gene regulation networks were developed as described. Briefly, we kept outgoing edges of expressed TFs from the union of high confidence general GRNs deposited in TRRUST (Han et al., 2018), RegNetwork (Liu et al., 2015), and DoRothEA (Garcia-Alonso et al., 2019; Holland et al., 2020), to convert this general network into context-specific networks. Improbable regulations were filtered by keeping those that arise from TF only if the TF is expressed as previously described (Martin et al., 2016; Santander et al., 2018). Additionally, we have integrated the information related to gene expression and promoter/CRE acetylation status. The Cytoscape session that includes these networks can be found in the supplementary data.

### Master Regulators

Master Regulators (MRs) search in our contextualized networks was performed as follows. For genes defined as differentially expressed, we select their first and second upstream neighbors in both networks similarly (Santander et al., 2018). Then, we looked for nodes with the highest edge density according to (Davis & Rebay, 2017). To do so, an iterative step deleting nodes with lower out-degree was performed until the deletion of a node in the subnetwork resulted in an unconnected network, with remaining TFs being categorized as candidates to be MR.

To determine if these candidates fulfill the definition of MR (Davis & Rebay, 2017), we have observed for candidate’s TFs that regulate the candidate, if its candidate is regulated by other candidates, and if the candidates have a physical interaction annotation among them (Szklarczyk et al., 2019).

## Declarations

### Ethics approval and consent to participate

The bioethics committee of Universidad Mayor, Chile, approved the protocol for animal management for this study.

### Consent for publication

The Author hereby consents to the publication of this work.

### Availability of data and materials

All sequence data are accessible with accession number BioProject ID: PRJNA873204 https://dataview.ncbi.nlm.nih.gov/object/PRJNA873204?reviewer=q2jn8g48s5gjf7hlo6b7a13h02

### Competing interests

The authors have declared that no competing interests exist.

### Funding

Beca de Doctorado Nacional ANID 21181038 and Beca Universidad Mayor (to S.A). ANID/CONICYT - FONDECYT Postdoctorado 3160622 (to M.S.) and 3220673 (to J.S.C.R). Proyecto Fondecyt regular 1181089 (to AM) and Fondecyt regular 1191003 (R.L.V.). Proyecto Fondecyt Inicio 11171015 (to M.A.S). Centro Ciencia & Vida, FB210008, Financiamiento Basal para Centros Científicos y Tecnológicos de Excelencia de ANID (to A.M). Powered@NLHPC: this research was partially supported by the supercomputing infrastructure of the NLHPC (ECM-02) (to A.M) and the computing infrastructure of the Centro de Genómica y Bioinformática, Universidad Mayor.

### Author Contributions

Formulation of research objectives and goals: S.A, A.M, and M.A.S.; Data curation: S.A.; Conducting biological experiments: S.A., M.S, M.C.O., R.L.V., and M.A.S; Conducting bioinformatic experiments: S.A.; Advice on bioinformatic analysis: J.S.R., A.M, and M.A.S; Editorial – original draft: S.A., J.S.C.R., R.L.V., A.M, and M.A.S; Proofreading and editing: S.A., J.S.C.R., R.L.V., A.M, and M.A.S. All authors read and approved the final manuscript.

## Acknowledgements

Not applicable

## Supplementary figures

**Supplementary figure 1:**
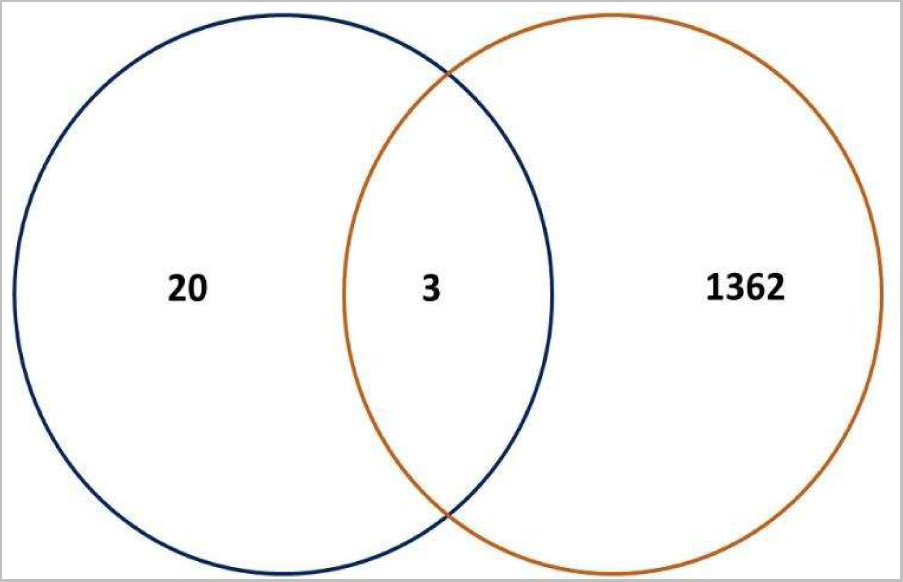
Venn diagram representation of differentially expressed genes in the 4-week and 6-week model R6/2 and control samples. It’s be seen in the union that 3 differentially expressed genes are present in the 2 analyzed ages.

**Supplementary figure 2:**
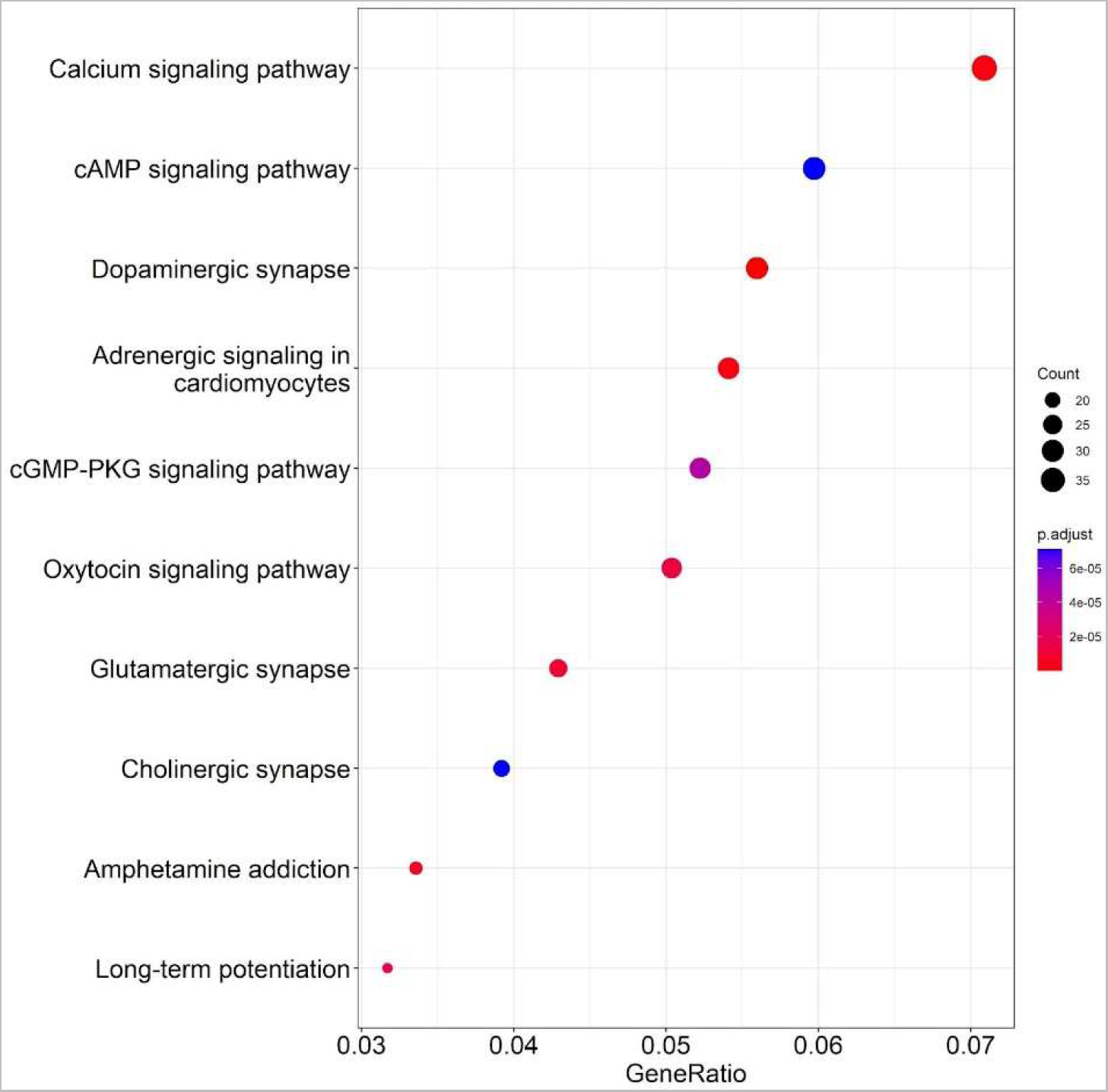
DE genes and Top 10 enrichment of metabolic pathways enriched in the R 6/2 model at 6 weeks.

**Supplementary figure 3:**
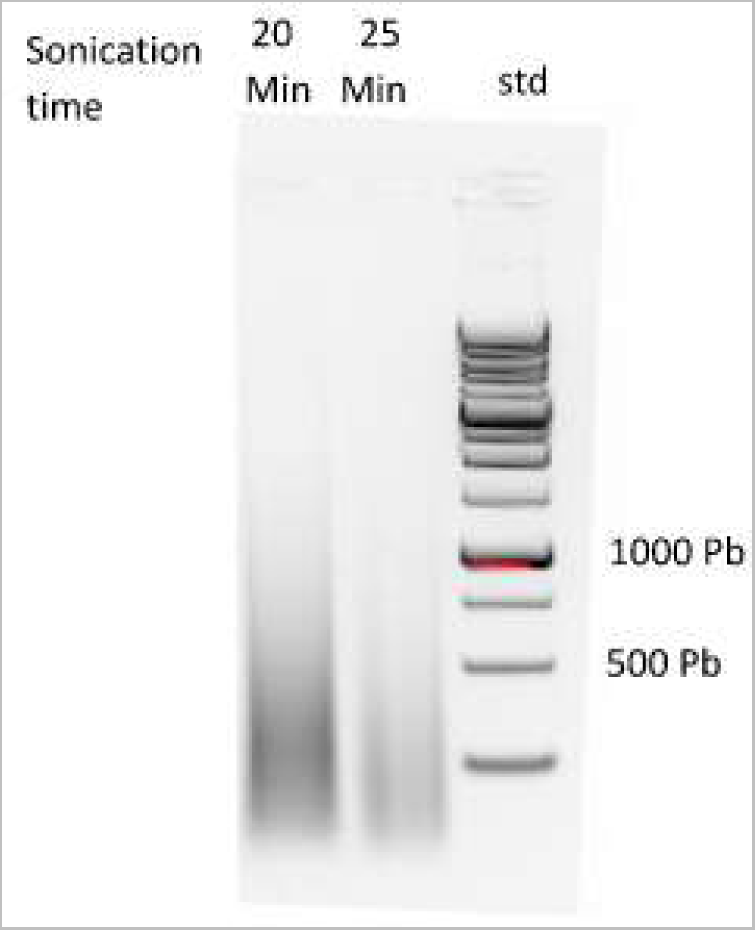
Gel electrophoresis image where chromatin fragmentation is observed when subjected to 20 and 25 minutes of sonication. Both bands are between 350 and 500 Pb.

**Supplementary figure 4:**
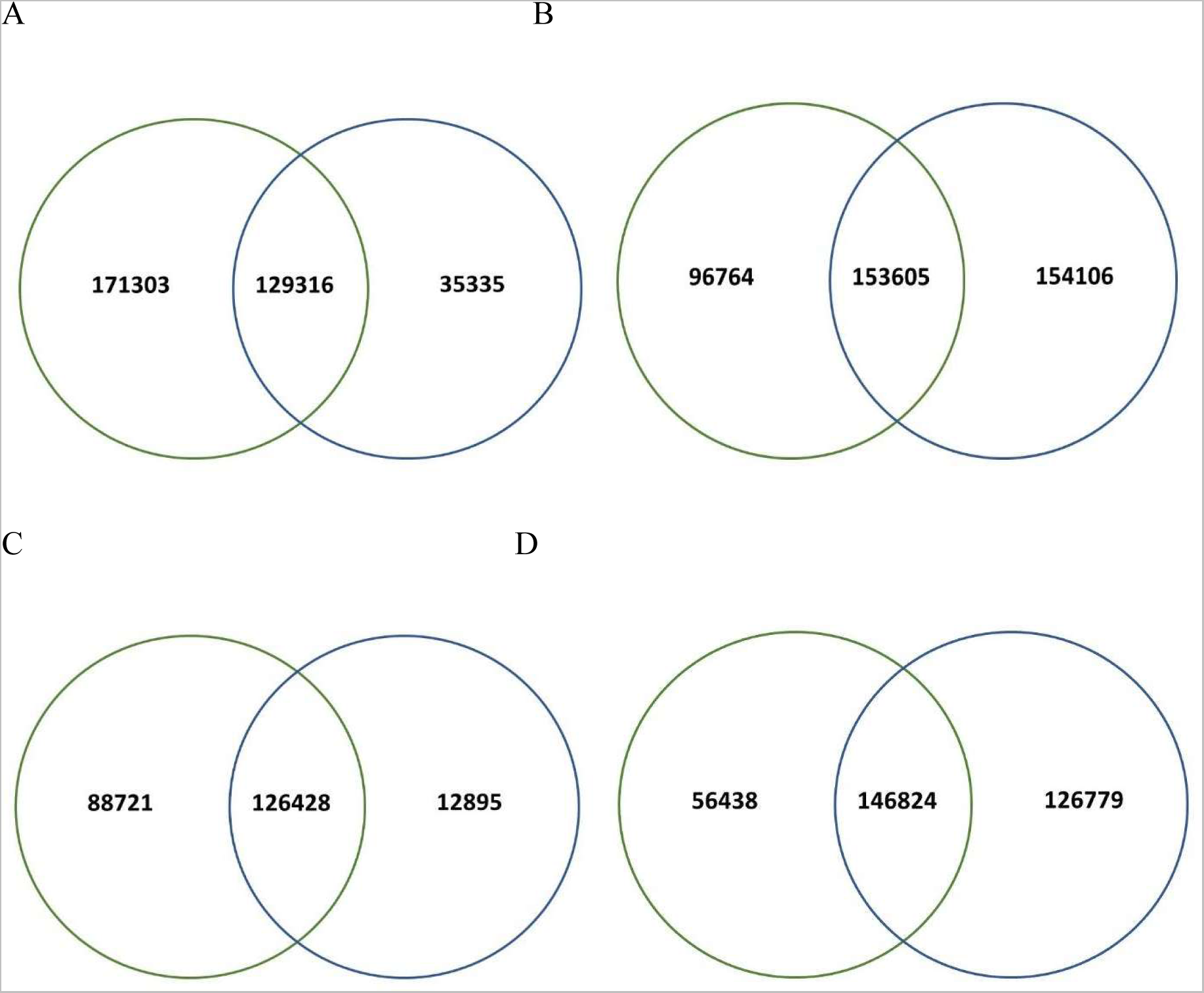
Venn diagrams for shared acetylation peaks in the control and mutant conditions at 4 and 6 weeks are represented using the bedtools software. The red circle represents the first replica, and blue color for the second replica of the ChIP-seq analysis in each of the experiments carried out. **A)** Venn diagram representation of the 4-week samples, there are 129316 peaks in common in the control samples. **B)** Representation of Venn diagram showing the samples R 6/2 4 weeks results, finding 153605 peaks in common. **C)** Representation of a Venn diagram showing the results of the 6-week control samples, finding 126428 peaks in common. **D)** Representation of Venn diagram showing the samples R 6/2 6-weeks results, finding 146824 peaks in common.

**Supplementary figure 5:**
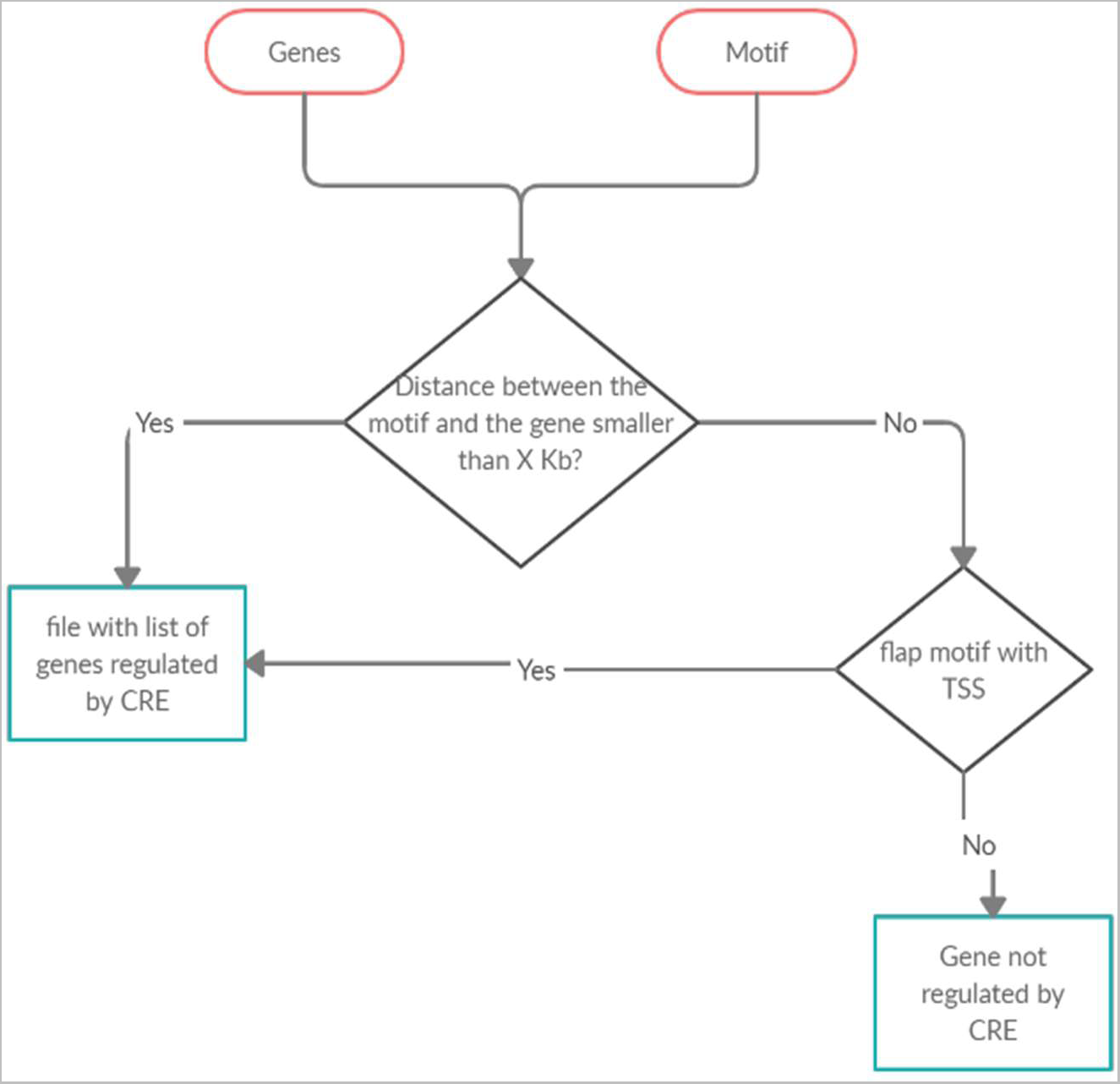
Flow diagram for the script that associate CRE regions where CREB could bind and regulate genes, proposing in this way candidate regulations.

**Supplementary figure 6:**
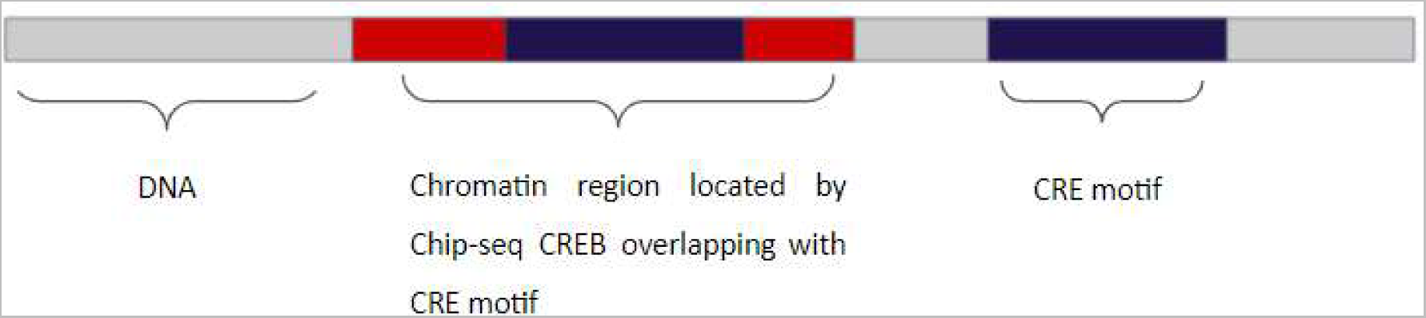
Validation scheme of CRE motif occurrences. The gray color represents the DNA. The red color corresponds to the chromatin region located by the Chip-seq CREB overlapping CRE motif. Then the blue color represents the CRE motif found by the script.

**Supplementary Table 1:** List of 81 genes validated and described in the literature is characterized as CRE-regulated genes.

|  |  |  |  |  |  |  |
| --- | --- | --- | --- | --- | --- | --- |
| *Nr4a2* | *Penk* | *Gpr3* | *Cdkn1a* | *Layn* | *Chrna1* | *Ociad2* |
| *Crhbp* | *Cd68* | *Tpm4* | *Ptprn* | *Btg2* | *Inhba* | *Rasgef1b* |
| *Fos* | *Ptgs2* | *Nfil3* | *Npas4* | *Fam150b* | *Ecm1* | *Homer1* |
| *Pcdh11x* | *Slc17a6* | *Cgref1* | *Scg2* | *Serinc2* | *Ier2* | *Gpr123* |
| *Nr4a1* | *Nap1l5* | *Gadd45g* | *Cyfip2* | *Ostf1* | *Peg10* | *Esd* |
| *Tinf2* | *Cnn1* | *Emd* | *Srxn1* | *Ifi202b* | *Akap13* | *Sat1* |
| *Dusp1* | *Tac1* | *Gadd45b* | *Sst* | *Spred3* | *Bag3* | *Ier3* |
| *Vgf* | *Ecel1* | *Sik3* | *S1pr3* | *Stat3* | *Tmbim1* | *Gadd45a* |
| *Fosb* | *Npy* | *Per1* | *Htr2c* | *Nrsn1* | *Cd9* | *Tagln2* |
| *Nptx2* | *Sik1* | *Gfra1* | *Fn1* | *Nnat* | *Vim* |  |
| *Crem* | *Sgk1* | *Synj2* | *Pvr* | *Stc1* | *Rfx4* |  |
| *Plat* | *Siah3* | *Cartpt* | *Litaf* | *Gem* | *Fgfr2* |  |

**Supplementary figure 7:**
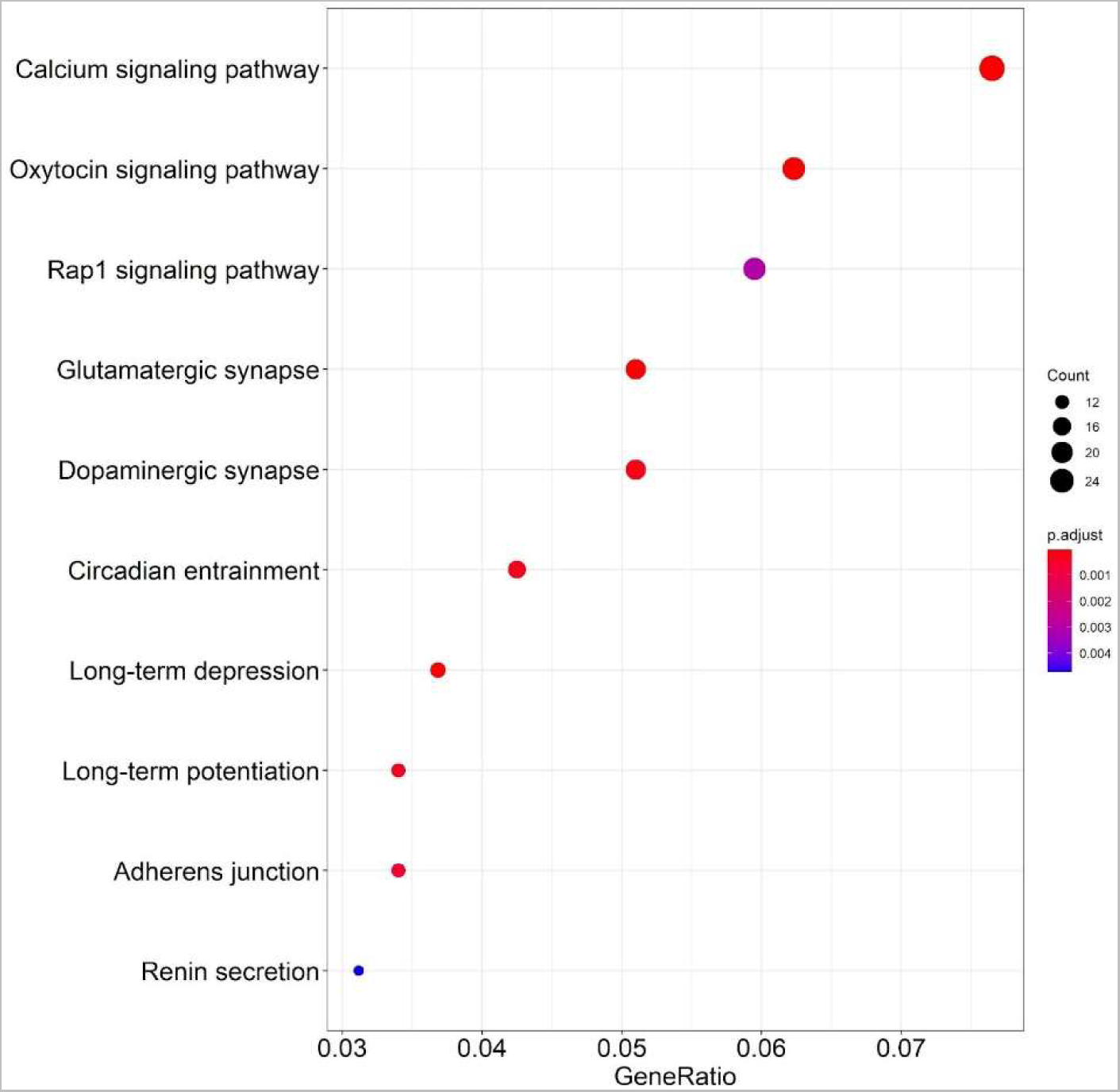
Distribution of differentially acetylated CRE regions at 6 weeks, in the Top 10 KEGG pathway analysis of differentially acetylated CRE regions.

**Supplementary Figure 8:**
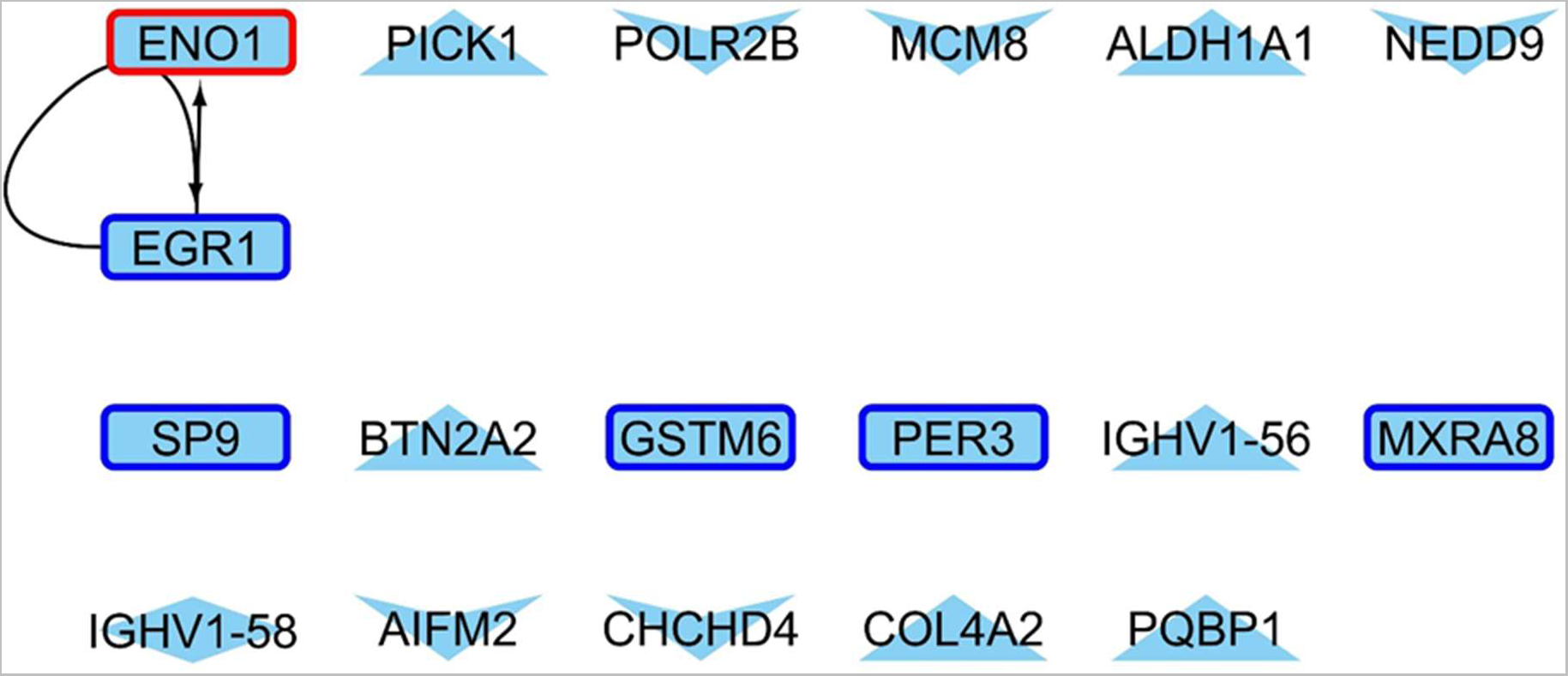
Genes with up (red border) or downregulated (blue) expression, CRE-region with increased acetylation pattern (diamond shape) and whose promoters present differential acetylation levels (triangle for up acetylation and V for low acetylation levels) 4 weeks old HD model.

**Supplementary Figure 9:**
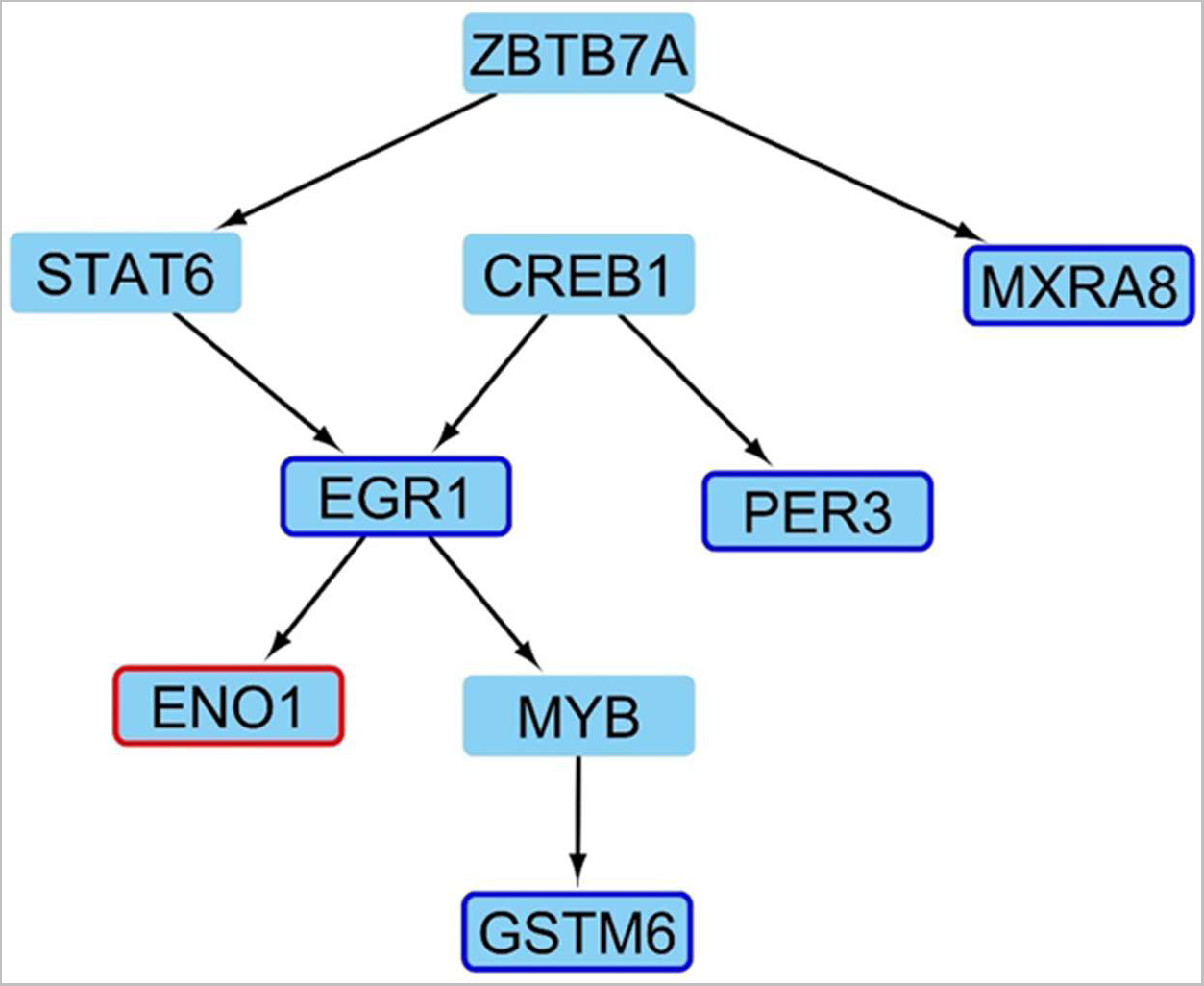
Candidates to be master regulators up (red borders) and down regulated (blue border) in HD condition and nodes that direct at 4 weeks age mice.

**Supplementary Figure 10:**
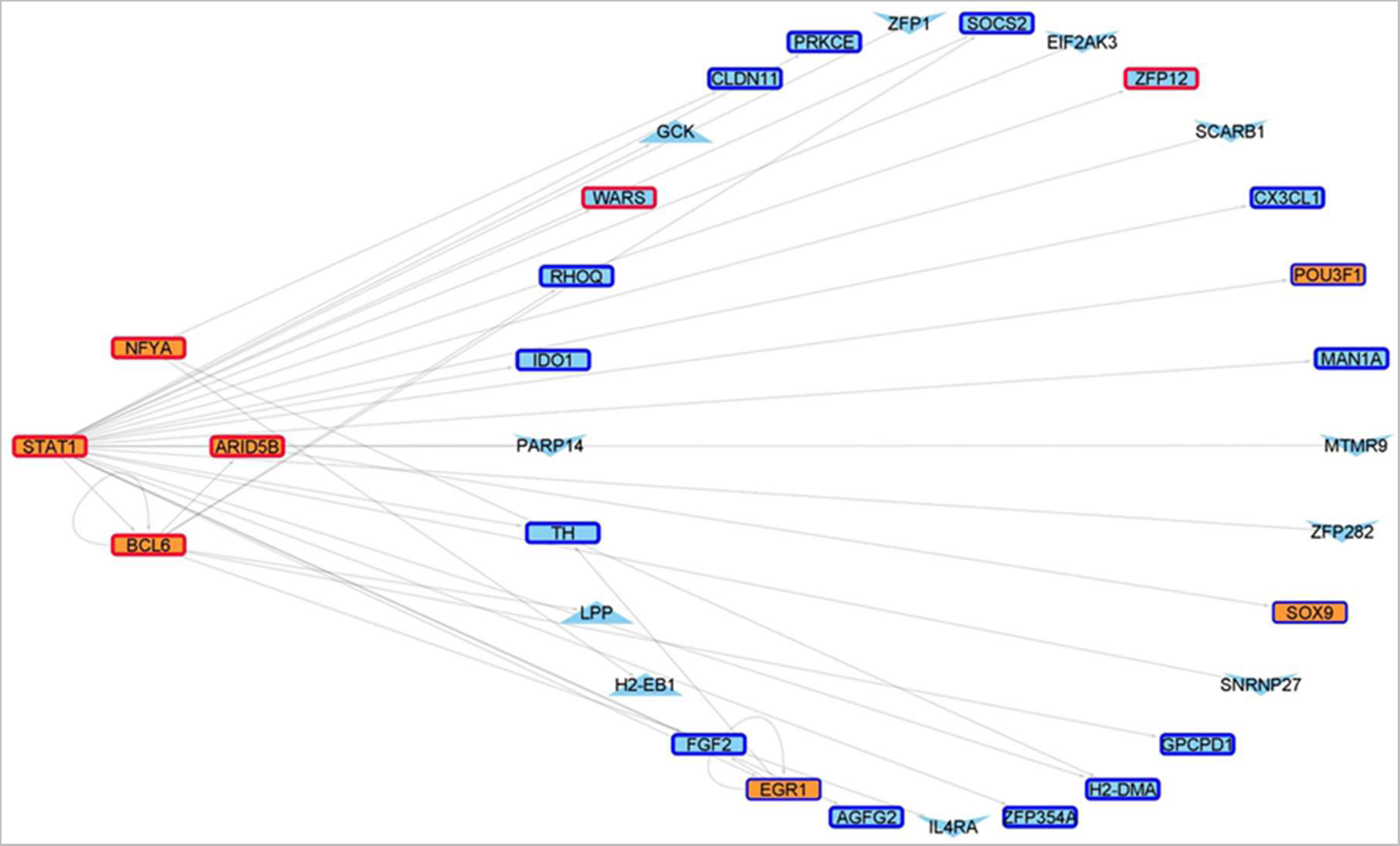
Master regulators (orange nodes) whose expression is upregulated (red border) in R 6/2 mice at 6 weeks old and genes that direct with changes in their expression levels (blue border: genes downregulated) or changes in their acetylation patterns (Triangles and V-shaped nodes represent genes whose promoters are up/down acetylated).

**Supplementary Figure 11:**
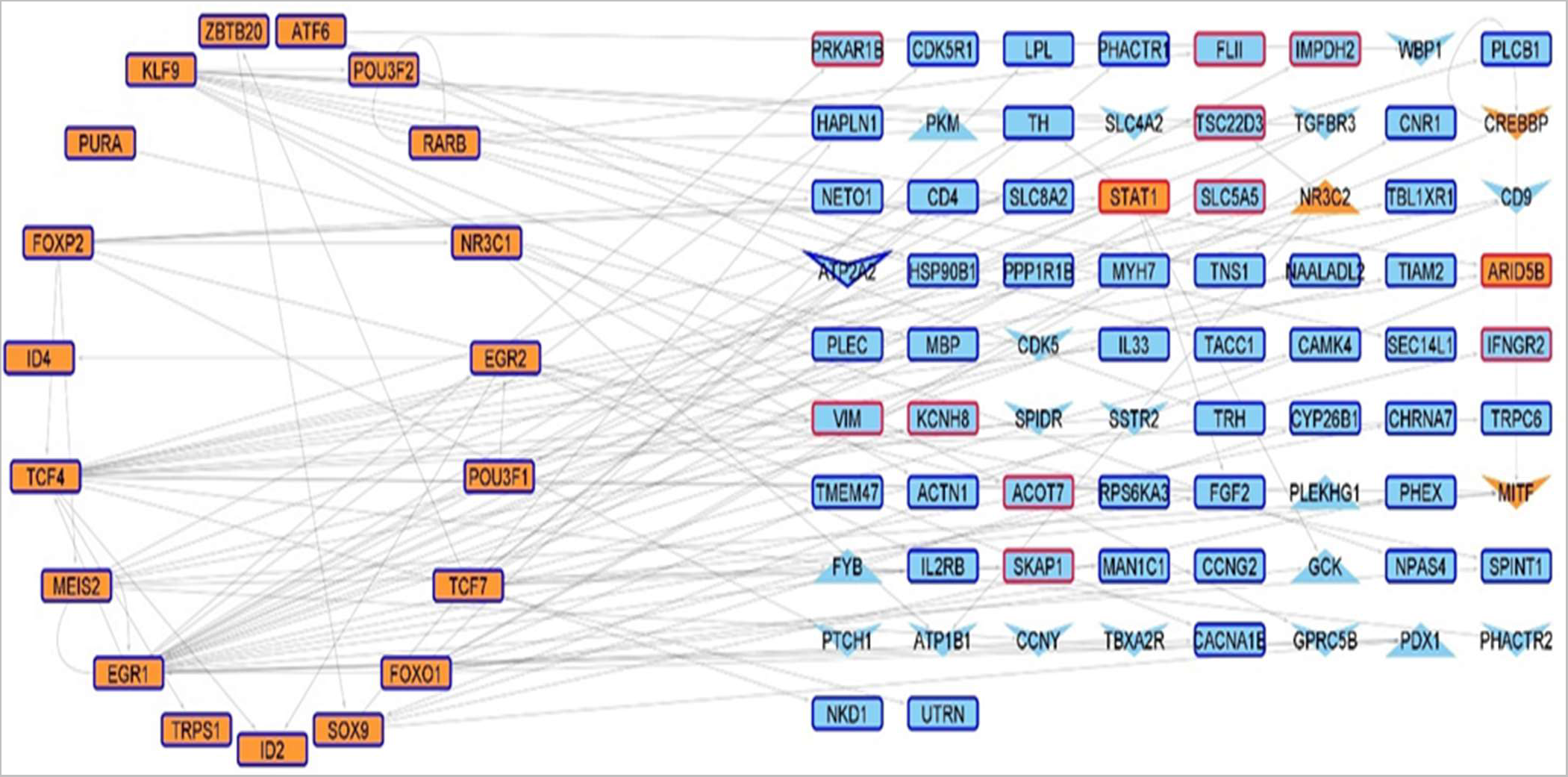
Master regulators (orange nodes) whose expression is downregulated (blue border) in R 6/2 mice at 6 weeks old and genes that direct with changes in their expression levels (red border: genes upregulated) or changes in their acetylation patterns. Triangles and V-shaped nodes represent genes whose promoters are up/down acetylated, whilst hexagons and diamonds represent genes with a CRE region up/down acetylated.

## Notes

### Competing Interest Statement

The authors have declared no competing interest.

