## Supplementary material for "Transcriptional and Histone acetylation changes associated with CRE elements expose key factors governing the regulatory circuit in early stage of Huntington’s disease models": Supp. Material: readme.txt

This file describe each of the supplemental files described in the manuscript.

DE_genes_4_weeks.csv : This file is the resulting table of DESeq2 for the differentially expressed (DEG) comparing mutant vs control condition at 4-week-old mice in the study. 
DE_genes_6_weeks.csv : This file is the resulting table of DESeq2 for the DEG comparing mutant vs control condition at 6-week-old mice in the study. 

DE_ac_genes_in_promoter_4_weeks.csv : This file is the resulting table of DESeq2 for promoters with differentially acetylated regions at 4-week-old mice. 
DE_ac_genes_in_promoter_6_weeks.csv : This file is the resulting table of DESeq2 for promoters with differentially acetylated regions at 6-week-old mice.

DE_ac_CRE_region_in_4_weeks.csv : This file is the resulting table of DESeq2 for of the CRE regions that are differentially acetylated in 4-week-old mice.
DE_ac_CRE_region_in_6_weeks.csv : This file is the resulting table of DESeq2 for of the CRE regions that are differentially acetylated in 6-week-old mice.

Cytoscape_session_4weeks.cys :This file correspond to the Cytoscape session contains the networks for control and mutant condition (and their node/edge selection) at 4-week-old mice in the study. 
Cytoscape_session_6weeks.cys :This file correspond to the Cytoscape session contains the networks for control and mutant condition (and their node/edge selection) at 6-week-old mice in the study. 
